# Ancient DNA reveals interstadials as a driver of the common vole population dynamics during the last glacial period

**DOI:** 10.1101/2022.08.09.503016

**Authors:** Mateusz Baca, Danijela Popović, Anna Lemanik, Sandra Bañuls-Cardona, Nicholas J. Conard, Gloria Cuenca-Bescós, Emmanuel Desclaux, Helen Fewlass, Jesus T. Garcia, Tereza Hadravova, Gerald Heckel, Ivan Horáček, Monika Vlasta Knul, Loïc Lebreton, Juan Manuel López-García, Eliza Luzi, Zoran Marković, Jadranka Mauch Lenardić, Xabier Murelaga, Pierre Noiret, Alexandru Petculescu, Vasil Popov, Sara E. Rhodes, Bogdan Ridush, Aurélien Royer, John R. Stewart, Joanna Stojak, Sahra Talamo, Xuejing Wang, Jan M. Wójcik, Adam Nadachowski

## Abstract

1

**Aim:** The common vole is a temperate rodent widespread across Europe. It was also one of the most abundant small mammal species throughout the Late Pleistocene. Phylogeographic studies of its extant populations suggested the Last Glacial Maximum (LGM, 26.5–19 ka ago) as one of the main drivers of the species’ population dynamics. However, analyses based solely on extant genetic diversity may not recover the full complexity of past population history. The main aim of this study was to investigate the evolutionary history and identify the main drivers of the common vole population dynamics during the Late Pleistocene.

**Location:** Europe

**Taxon:** Common vole (*Microtus arvalis*)

**Methods:** We generated a dataset comprising 4.2 kb-long fragment of mitochondrial DNA from 148 ancient and 51 modern specimens sampled from multiple localities across Europe and covering the last 60 thousand years (ka). We used Bayesian inference to reconstruct their phylogenetic relationships and to estimate the age of specimens that were not directly dated.

**Results:** We estimate the time to the most recent common ancestor of all Last Glacial and extant common vole lineages to 90 ka ago and the divergence of the main mtDNA lineages present in extant populations to between 55 and 40 ka ago, earlier than previous estimates. We find multiple lineage turnovers in Europe in the period of high climate variability at the end of Marine Isotope Stage 3 (MIS 3; 57–29 ka ago) in addition to those found previously around the Pleistocene/Holocene transition. Conversely, data from the Western Carpathians suggest continuity throughout the LGM even at high latitudes.

**Main conclusions:** Our results suggest that the main factor affecting the common vole populations during the last glacial period was the reduction of open habitats during the interstadial periods while the climate deterioration during the LGM had little impact on species’ population dynamics.

## 3 Introduction

The climatic and environmental changes during the last glacial period (ca. 115–11.7 ka ago) had a great impact on the evolutionary histories of most species. It has been suggested that species have responded to those changes according to their individual characteristics and there is no basis for considering them as communities responding to climate and environmental changes in the same manner (Baca et al., 2017; Lorenzen et al., 2011; Stewart, Lister, Barnes, & Dalén, 2010). However, within the ecosystem, species depend on a whole range of interactions at different trophic levels and across different ecological niches (Walther, 2010). Thus, investigations of species with different adaptations can reveal the spectrum of responses to the same climatic and environmental fluctuations and allow identification of the key factors driving ecosystem responses (Cooper et al., 2015). Small mammals may be especially well suited for such investigations as, in contrast to megafaunal species, they seem to be little affected by activities of Palaeolithic hunter-gatherers and their population dynamics were mainly driven by environmental changes.

The common vole *Microtus arvalis* (Pallas, 1779) is a temperate rodent species which is present in most of continental Europe (Figure 1b). The species feeds primarily on leaves and grasses and prefers well-drained grasslands, pastures, and alpine meadows from lowlands up to ca. 3000 m above sea level. At present it mainly utilizes secondary habitats, such as agricultural fields, where it is often considered a pest (Jacob, Manson, Barfknecht, & Fredricks, 2014). The earliest remains of ancestral common voles (*Microtus arvalis*-group) in Europe date to ca. 0.6-0.5 Mya (Berto, Nadachowski, Pereswiet-Soltan, Lemanik, & Kot, 2021; Kučera, Suvova, & Horáček, 2009; Maul & Markova, 2007). The fossil record from the last glacial period (115– 11.7 ka ago), attests to its continuous presence on most of the continent (Chaline, 1972; Horáček & Ložek, 1988; Jánossy, 1986; Nadachowski, 1989). In many localities on the European Plain, from France to Poland, the common vole was the most abundant small mammal, alongside the collared lemming (*Dicrostonyx torquatus*) and the European narrow-headed vole (*Lasiopodomys anglicus*) (Royer et al., 2016; Socha, 2014). The mitochondrial DNA (mtDNA) phylogeography of extant populations of the common vole has been intensively studied, with the aim of reconstructing the post-glacial history of the species. The extant mtDNA diversity is partitioned into six divergent lineages with parapatric distribution: Western-South (WS), Western-North (WN), Italian (ITA), Balkan (B), Central (CEN), and Eastern (E) (Bužan, Förster, Searle, & Kryštufek, 2010; Haynes, Jaarola, & Searle, 2003; Heckel, Burri, Fink, Desmet, & Excoffier, 2005; Stojak, McDevitt, Herman, Searle, & Wójcik, 2015) (Figure 1b). Most of the previous studies estimated the time to the most recent common ancestor (tMRCA) of the extant common vole populations to between 65 and 50 ka ago and subsequent divergence of main lineages to between 50 and 20 ka ago (García et al., 2020; Heckel et al., 2005; Stojak et al., 2016). In contrast, Fink et al. (2004) and Tougard et al. (2008) used fossil calibration to suggest a much older diversification in the Middle Pleistocene. Analyses of nuclear DNA revealed overall a very good correspondence with the spatial distributions of mtDNA lineages (Fischer, Foll, Heckel, & Excoffier, 2014; Heckel et al., 2005) and detailed analyses of the contact areas between lineages demonstrated admixture only in narrow hybrid zones (Beysard & Heckel, 2014; Braaker & Heckel, 2009). However, the divergence time estimates of the evolutionary lineages based on the nuclear data were generally much more recent than those based on mtDNA and suggested that the diversification of common vole evolutionary lineages took place during or after the LGM (Heckel et al., 2005; Lischer, Excoffier, & Heckel, 2014).

**Figure 1.**
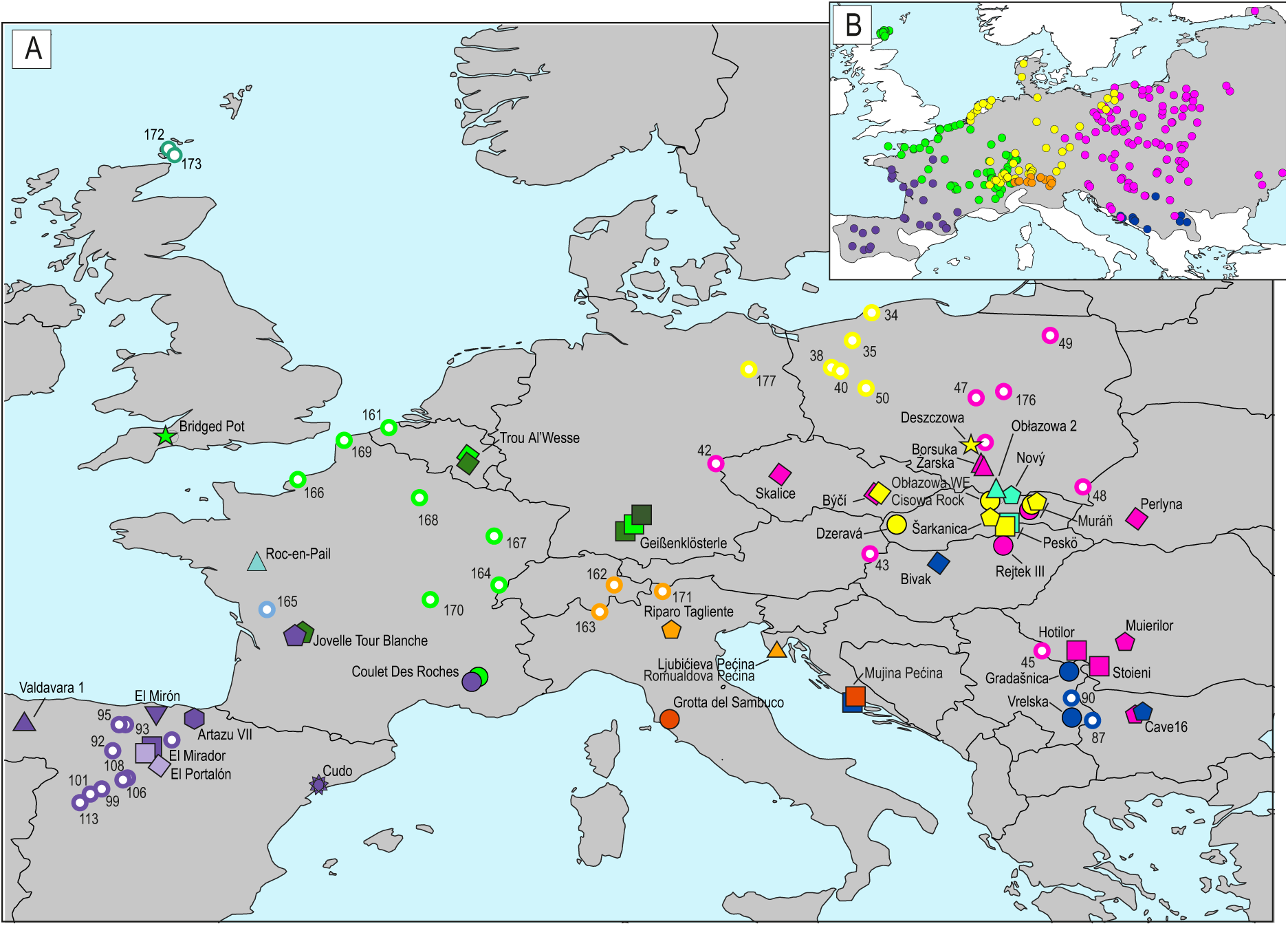
Sampling localities of modern and ancient common voles across Europe (a). Filled symbols represent paleontological sites, while unfilled circles denote localities of modern specimens. Numbers near modern localities correspond to specimen numbers (WM…) in Figure 2 and Table S2. Symbols and names are consistent with those in Figure 2 and are coloured according to the vole mtDNA lineage found at the site. If multiple mtDNA lineages were found at the site, two or more symbols are presented. Distribution of the main mtDNA lineages in extant populations (b). The grey area depicts the current range of common vole in Europe. The coloured circles represent sampling localities of the common vole and mtDNA lineages compiled from previous studies: pink – Eastern (E); yellow – Central (CEN); orange – Italian (ITA); green – Western-North (WN); violet – Western-South (WS); navy blue – Balkan (B).

The present-day distribution of common vole mtDNA lineages (Figure 1b) was interpreted as evidence for the survival of common vole populations during the LGM in both traditional Southern glacial peninsular refugia, as well as at higher latitudes, especially in Central France (WN), north of the Alpine region (CEN) and in the Carpathian area (E) (Heckel et al., 2005; Stojak et al., 2015; Tougard et al., 2008). Examination of nuclear DNA from multiple European populations revealed a south-west cline of genetic diversity which was interpreted as indicative of a westward expansion of the common vole at a time prior to the LGM (Heckel et al., 2005), nevertheless the pre-LGM history of the species remains largely unknown.

A detailed study of the Eastern mtDNA lineage suggested that it originated in the Carpathian area and its current distribution is the result of an expansion which started after the LGM with a possible bottleneck during the Younger Dryas (12.8–11.7 ka ago) (Stojak et al., 2016). Recently, an ancient DNA investigation of the common vole remains from the Western Carpathians has shown the presence of the Eastern lineage from the early Holocene onwards. However, it also showed that from at least 18 ka ago the Central lineage was present in this region suggesting a population replacement around the Pleistocene/Holocene transition (Baca et al., 2020). This challenged the simple model of post-glacial recolonisation of the Eastern lineage from the Carpathian refugium and suggested that the analyses based solely on extant genetic diversity may not recover the full complexity of the Late Pleistocene population dynamics. To refine the evolutionary history of common vole populations in Europe and to compare it with the past histories of coeval cold-adapted species such as the collared lemming, we generated and analysed a new mitochondrial dataset consisting of nearly 200 sequences from Late Pleistocene and extant individuals.

## 4 Material and Methods

### 4.1 Ancient specimens

Isolated lower first molars or mandible fragments with molars classified as *M. arvalis* or *Microtus sp*. based on morphology of the occlusal surface were collected from various palaeontological sites across Europe (Table S1). Each tooth was photographed at the Institute of Systematics and Evolution of Animals, PAS.

### 4.2 Modern specimens

DNA of modern specimens from various locations across Europe was extracted previously (Table S2). A target 4.2 kb region of mtDNA spanning positions 12,000 to 16,247 according to the reference sequence (NC_038176; Folkertsma et al., 2018) was generated using various approaches. It was either PCR amplified, sonicated, transformed into sequencing libraries and sequenced or the genomic DNA was sonicated and transformed into sequencing libraries. It was then either enriched for target fragment using in-solution hybridization and sequenced or the target region was extracted from deep shotgun sequencing data (see Appendix A1.1 and Table S2 for more details).

### 4.3 Ancient DNA extraction, target enrichment and sequencing

DNA extraction and pre-PCR library preparation steps were performed in the dedicated ancient DNA laboratory at the Centre of New Technologies at the University of Warsaw. Each tooth was thoroughly cleaned with ultra-pure water in a 2 ml tube and crushed with a pipette tip. DNA was extracted following a silica spin column-based protocol optimized for retrieval of short DNA molecules (Dabney et al., 2013). A negative control without biological material was processed alongside each batch of 15 specimens. Double-stranded, double-indexed sequencing libraries were produced from half of the DNA extract (20 µl) following a previously established protocol (Meyer & Kircher, 2010) with minor modifications (Baca et al., 2019). For some specimens that yielded low-quantity DNA additional double-indexed, single-stranded sequencing libraries were prepared following the protocol proposed by Gansauge et al. (2020) (see Appendix A1.3–A1.4 for more details).

Libraries were enriched for vole mitochondrial DNA using an in-solution hybridization protocol described in Baca et al. (2019). Up to five libraries were pooled for hybridization reaction. We performed two rounds of hybridization in 65ºC for 22–24h each. After each round, library pools were washed and amplified in triplicate for 10 to 15 cycles. Enriched library pools were combined, quantified using qPCR and sequenced on Illumina NextSeq550 platform (MID output, 2×75 bp kit; see Appendix A1.2–A1.5 and Tables S4–S5 for more details).

### 4.4 Sequence processing

Sequencing reads were demultiplexed using bcl2fastq v. 2.19 (Illumina). Overlapping reads were collapsed, adaptor and quality trimmed using AdapterRemoval v. 2.2.2 (Schubert, Lindgreen, & Orlando, 2016). Then, reads were mapped to the common vole mtDNA genome using the *mem* algorithm in bwa v. 0.7.17 (Li & Durbin, 2010). Duplicates, short (<30 bp) and low mapping quality reads (mapq<30) were removed using *samtools* v. 1.9. Variants and consensus sequences were called using *bcftools* v. 1.9 (Li et al., 2009). Read alignments and vcf files were inspected manually using Tablet v. 1.17 (Milne et al., 2013). Positions with coverage below three were masked with N. If a base was supported by less than 75% of reads, an IUPAC symbol was inserted. MapDamage v. 2.08 (Jónsson, Ginolhac, Schubert, Johnson, & Orlando, 2013) was used to assess the damage patterns and length distribution of DNA molecules. See Appendix A1.6 for more details.

### 4.5 Phylogenetic analyses and molecular dating of specimens

We used a Bayesian approach, implemented in BEAST 1.10.4 (Suchard et al., 2018), to estimate divergence times of common vole lineages and the age of specimens which were not directly dated. For the phylogenetic inference we used only ancient and modern sequences with at least 70% and 90% of the target mtDNA fragment (4.2 kb) recovered, respectively. Sequences were aligned with MAFFT v. 7.407 (Katoh & Standley, 2013). The best substitution model selected by jModelTest2 (Darriba, Taboada, Doallo, & Posada, 2012), TIM2+F+I+G4, was not easily available in BEAST we therefore used the closest available one: GTR +I+G.

First, we used Bayesian evaluation of temporal signal (BETS) (Duchene et al., 2020) to check whether there is sufficient temporal resolution within our dataset to calibrate the molecular clock. We used all directly radiocarbon dated (n = 20) and modern (n = 51) specimens and tested four alternative models. In two of them, we assigned real sampling times to the sequences (heterochronous analysis) and used either strict clock or uncorrelated relaxed lognormal clock. In the two other models, we used the same sampling time for all sequences (isochronous analysis) and applied either strict clock or uncorrelated relaxed lognormal clock (see Appendix A1.7 for more details). Then, we performed the leave-one-out analysis on the directly radiocarbon dated specimens to check the accuracy of the age estimates produced using the available calibration dataset. In this analysis we estimated the age of each directly radiocarbon dated specimen using all the remaining radiocarbon dated and modern specimens to calibrate the molecular clock. Next, we estimated the age of each ancient, not directly dated, specimen (n = 128) in a separate BEAST run, again using all directly dated and modern specimens to calibrate the molecular clock. Finally, we ran a joint analysis with all the sequences. For the specimens that were not directly dated we set a lognormal prior with a mean equal to the mean age estimated in the individual analysis and the range covering the 95% HPD interval of the individual estimate (see Appendix A1.7, Tables S6–S9 for more details).

The demographic reconstructions using the Bayesian Skyline and Bayesian SkyGrid methods implemented in BEAST 1.10.4 were performed for the WN lineage specimens from Spain (n=58). In the case of other localities either the number of available sequences was low or the assumption of population continuity (i.e., lack of lineage turnovers) were violated (see Appendix A1.8 for details).

### 4.6 Radiocarbon dating

Selected vole mandibles were pretreated for radiocarbon dating in the Department of Human Evolution at the Max Planck Institute for Evolutionary Anthropology (MPI-EVA, Leipzig, Germany) following the protocol for <100 mg bone samples described in Fewlass et al. (2019). The quality of the collagen extracts was assessed based on the collagen yield as a percentage of the original bone weight (minimum requirement 1%). The elemental and isotopic ratios of the extracts (∼0.5 mg) were measured at the MPI-EVA on a Thermo Finnigan Flash elemental analyser coupled to a Thermo Delta plus XP isotope ratio mass spectrometer (EA-IRMS). Where sufficient collagen was extracted, collagen was graphitised using the automated graphitisation equipment (AGE) (Wacker, Němec, & Bourquin, 2010) in the Lab of Ion Beam Physics at ETH-Zurich (Switzerland) and dated on a MIni CArbon DAting System (MICADAS) accelerator mass spectrometer (AMS) (Wacker, Bonani, et al., 2010). Where the extracted collagen yield was insufficient for graphitization, it was combusted to CO_2_ and measured directly using the gas interface system coupled to the gas ion source of the MICADAS (Wacker et al., 2013) following the protocol described in Fewlass et al. (2019) (see Appendix A1.9 for more details).

To improve stratigraphic information available for the sites from which the analysed specimens originated we also obtained radiocarbon dates from five palaeontological sites (Appendix A1.10, Table S10). Radiocarbon dates were calibrated in OxCal v4.4 (Bronk Ramsey, 2009) using the IntCal20 (Reimer et al., 2020) calibration curve.

## 5 Results

We generated a dataset of 4.2 kb long mtDNA sequences from 199 ancient and modern common vole specimens (148 ancient and 51 modern). Sequences of 82 ancient specimens are reported for the first time and either 4.2 kb or 1kb fragments of mtDNA of the remaining 66 specimens were reported previously (Baca et al., 2020; Baca et al., 2021; Lemanik et al., 2020) (Table S1). All 82 newly reported specimens yielded short inserts and elevated level of cytosine deamination at the terminal nucleotides characteristic for ancient DNA (Table S1). Ancient specimens which yielded mtDNA sequences came from 40 sites scattered across Europe and covered the period of the last ca. 60 thousand years (Table S1, Figure 1a). MtDNA cytochrome *b* sequences of most modern specimens were reported previously (García et al., 2020; Stojak et al., 2016) and here only the longer mtDNA fragment was generated to increase the phylogenetic resolution. Direct radiocarbon dating was undertaken for 12 vole mandibles and 10 yielded collagen of sufficient quality for AMS dating (Table S3).The carbon to nitrogen ratio (C:N) of one specimen from Geißenklösterle (MI935) was at the limit (3.6) of accepted values for well-preserved collagen (2.9-3.6; Van Klinken, 1999) and yielded a relatively recent date with respect to the latest Electron Spin Resonance dating of the cave sediments (Richard et al., 2019) indicating that contamination with modern carbon, and therefore under-estimation of the true age, is likely. We therefore discarded this radiocarbon date and estimated the age of this specimen using the molecular approach. All other direct dates are regarded as reliable based on their chemical indicators (collagen yield, stable isotopic and elemental values; Table S3). As a result, we used sequences of 20 directly radiocarbon dated and 51 extant specimens to calibrate the molecular clock. The analysis of temporal signal (BETS) showed that the dated specimens included in our dataset are suitable for calibration of the molecular clock. The heterochronous model, with a strict clock and correct sampling times assigned to specimens, was strongly supported over all other models (2lnBF > 9) (Table S4). The leave-one-out analysis revealed that the dated dataset enables relatively accurate estimation of specimen ages, although in the case of three specimens (MI074, MI1337 and MI1355) the 95% highest posterior density intervals (95% HPD) of estimated ages did not overlap within 2-sigma ranges of the radiocarbon dates (Figure S1; Appendix A). In addition, the estimated ages of most specimens agreed with their stratigraphic position, providing evidence for accuracy of this approach (Table S1).

### 5.1 Diversification of common vole mtDNA lineages

The maximum clade credibility tree obtained in BEAST 1.10.4 (Figure 2), recovered, with high support, all six mtDNA lineages characterised previously in analyses of the mtDNA cytochrome *b* of modern individuals (Bužan et al., 2010; Haynes et al., 2003; Heckel et al., 2005). In addition, we identified three lineages which were only present in the Late Pleistocene specimens. These lineages were named by their geographic provenance and phylogenetic position: WNII and WNIII in Western Europe (Germany and France) and ITAII in Italy and Croatia (Figure 1a and 2).

**Figure 2.**
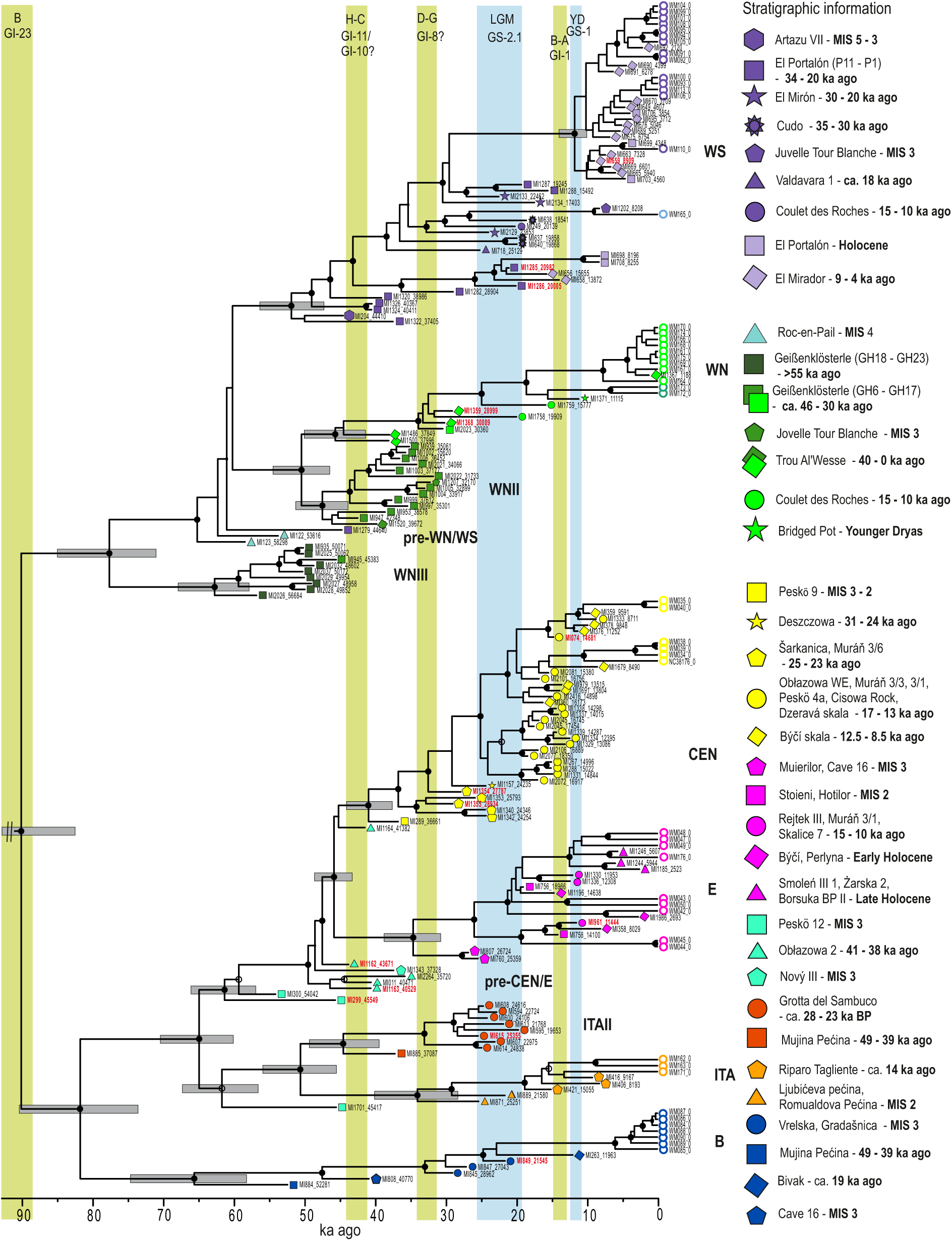
Maximum clade credibility tree of common vole mtDNA obtained in BEAST 1.10.4 and calibrated with radiocarbon dated specimens. Black dots indicate nodes with posterior probability above 0.95 and grey bars show the 95% HPD intervals of node ages. The tips are annotated with sample names, colour symbols (consistent with those in Figure 1) and, medians of age estimates (black font) or medians of calibrated radiocarbon dates (red font). The empty circles represent modern specimens. The legend on the right side of the figure provides information on stratigraphy and dating of the source sites and layers. The green and blue strips indicate the main interstadials and stadials: B – Brørup; H–C – Hengelo–Charbon; D–G Denekamp–Grand Bois; LGM – Last Glacial Maximum, B–A – Bølling– Allerød; YD – Younger Dryas.

The tMRCA of all European common voles was estimated to 90 ka ago (95% HPD: 98– 83 ka ago). Divergence time estimates of the subsequent lineages ranged between 82 ka ago (95% HPD: 91–75 ka ago) for the split between B and ITA/C/E, through 77 ka ago (95% HPD: 84–70 ka ago) for the split between WNIII and WNII/WN/WS to ca. 50–45 ka ago for the splits between a number of lineages: ITA and ITAII; CEN and E; WNII and WN (Figure 2).

### 5.2 Temporal population structure and dynamics of the common vole

#### 5.2.1 Western and Southwestern Europe

The oldest specimens from Western Europe were classified as belonging to the WNIII lineage which was the sister to the WS and WN/WNII lineages. The WNIII lineage contains mainly individuals from the lowermost layers of Geißenklösterle (GH23-18). The age of these individuals was estimated to between 56 and 45 ka (Figure 1a, 2, Table S1). Three haplotypes of similar age as the latter (57–44 ka ago) from Western France (Roc-en-Pail; MI122, MI123) and Northern Spain (El Portalon P9; MI1279) are located at the base of the WS, WNII and WN lineages (pre-WN/WS; Figure 1a, Table S1). Around 42 ka ago the WS lineage appeared in Northern Spain and the WNII lineage appeared in France, Belgium and Germany. The latter, composed of specimens from the younger layers of Geißenklösterle, Trou Al’Wesse and Jovelle, disappeared around 32 ka ago. The age of the oldest specimens from the WN lineage, which come from the Trou Al’Wesse, was estimated to 37 ka ago. This lineage was found among Late Pleistocene specimens from Western Germany, Belgium, France, and the UK.

The record from Spain suggests population continuity throughout the last 40 ka, although nearly all modern and Holocene specimens coalesce around 11.5 ka ago (Figure 2), which suggests significant reduction of population size around the Pleistocene/Holocene transition. This was further confirmed by the Bayesian demographic analysis, which suggest about five-fold reduction of female effective population size around the Pleistocene/Holocene transition with the minimum around the Early Holocene (11.7–9 ka ago), followed by a slight recovery around the Middle Holocene (9–6 ka ago) (Appendix A1.8, Figures S2 and S3). The WS lineage was also detected in southern France at Jovelle and Coulet des Roches during MIS 2 (29–14 ka ago).

#### 5.2.2 Central and Southeastern Europe

The oldest specimens in Central and Southeastern Europe come from Obłazowa 2, Nový and layer 12 of Peskö, all from the Western Carpathians. Their ages were estimated to between 55 and 35 ka ago and they hold a basal position with respect to Central and Eastern lineages (pre-CEN/E). The two oldest specimens from Peskö (MI299, MI300) were noticeably more divergent than the others. Starting from 35 ka ago we found the CEN lineage in the Western Carpathians as well as to their north, and the most recent specimens belonging to this lineage were dated to the Early Holocene. The earliest specimens assigned to the Eastern lineage were estimated to ca. 27–25 ka ago and were found in Central Romania (Muierilor Cave; MI760) and Northwestern Bulgaria (Cave 16; MI807). The more recent specimens from this lineage seem to represent a north and northwestern expansion of this population, with the first appearance in the Ukrainian Carpathians in Perlyna at around 14 ka ago and in the Western Carpathians around 12 ka ago (Rejtek III, Muráň 3/1).

The Balkan lineage occupied the same area as its current distribution starting from at least 50 ka ago (Mujina pećina). A single specimen from Bivak Cave in the Northern Hungary suggests that prior to the Holocene, its range extended further to the northeast.

The ITA and ITAII lineages were detected on the Italian and Balkan Peninsulas. ITAII specimens were found in central Italy (Grotta del Sambuco) and in middle Dalmatia (Mujina pećina) with ages estimated to between 36.5 and 18.5 ka ago. These were generally older than specimens from the ITA lineage, dated to 22.6–0 ka ago and found in northern Italy (Riparo Tagliente) and Istria (Ljubićeva pećina). The single individual from layer 12 of Peskö cave (Western Carpathians; MI1701), directly radiocarbon dated to 45 ka cal BP, was located at the base of the ITA lineages.

## 6 Discussion

The mitochondrial diversity of the common vole has been studied in detail to reconstruct the evolutionary history of species and elucidate its reactions to climate change. However, inferences from modern mitochondrial diversity are limited, since the signal of the population history is usually erased by the most recent reduction of effective population size (Mourier et al., 2012). Ancient DNA enables direct observation of changes in genetic diversity in response to climate or environmental changes by sampling populations prior and after such changes. Another great advantage of ancient DNA is that directly dated specimens may be used to estimate substitution rates and to calibrate the molecular clock providing robust estimates of divergence times without the need of external calibration. In this study, we make use of these advantages and reconstructed the evolutionary history of the common vole at much greater temporal depth providing new insights into its paleoecology.

### 6.1 The effects of climatic changes on diversification of common vole lineages

The estimated tMRCA of mtDNA for the European common vole (90 ka ago; 95% HPD: 98–83 ka ago) is substantially older than some previous estimates (García et al., 2020; Heckel et al., 2005; Stojak et al., 2015). It was similar to the recent estimates for the initial diversification of mtDNA lineages of the cold-adapted collared lemmings (100 ka ago, 95% HPD: 109–92 ka ago; E. Lord, personal communication). It was suggested that this may be an effect of a bottleneck during the Eemian Interglacial (MIS5e). It may also be related to the Brørup Interstadial (MIS 5c; ∼GS-23; ca. 104–88 ka). Vegetation during the Brørup Interstadial was characterised by temperate deciduous forests in Western Europe and boreal forests more to the north and east (Guiter et al., 2003; Helmens, 2014), providing unfavourable habitats for the cold-adapted collared lemming as well as for the common vole, both of which rely on various types of open habitat.

The initial divergence of the common vole lineages may have been the result of survival in two distinct refugial areas in the Alpine and Carpathian regions causing a partition in the mitochondrial diversity of the species in with two geographical areas. Western Europe, including the territories of present-day Spain, France, Switzerland, and parts of Germany, was occupied by the WNIII, WNII, WN and WS lineages. Meanwhile, Central and South-eastern Europe, including territories to the East and to the South of Germany, was occupied by the CEN, ITAII, ITA, B and E lineages. This geographic partitioning was maintained for at least 45 ka (Figure 3) and probably contributed to patterns of restricted geneflow and partial reproductive isolation of the present-day populations of Western-North and Central lineages (Beysard & Heckel, 2014).

**Figure 3.**
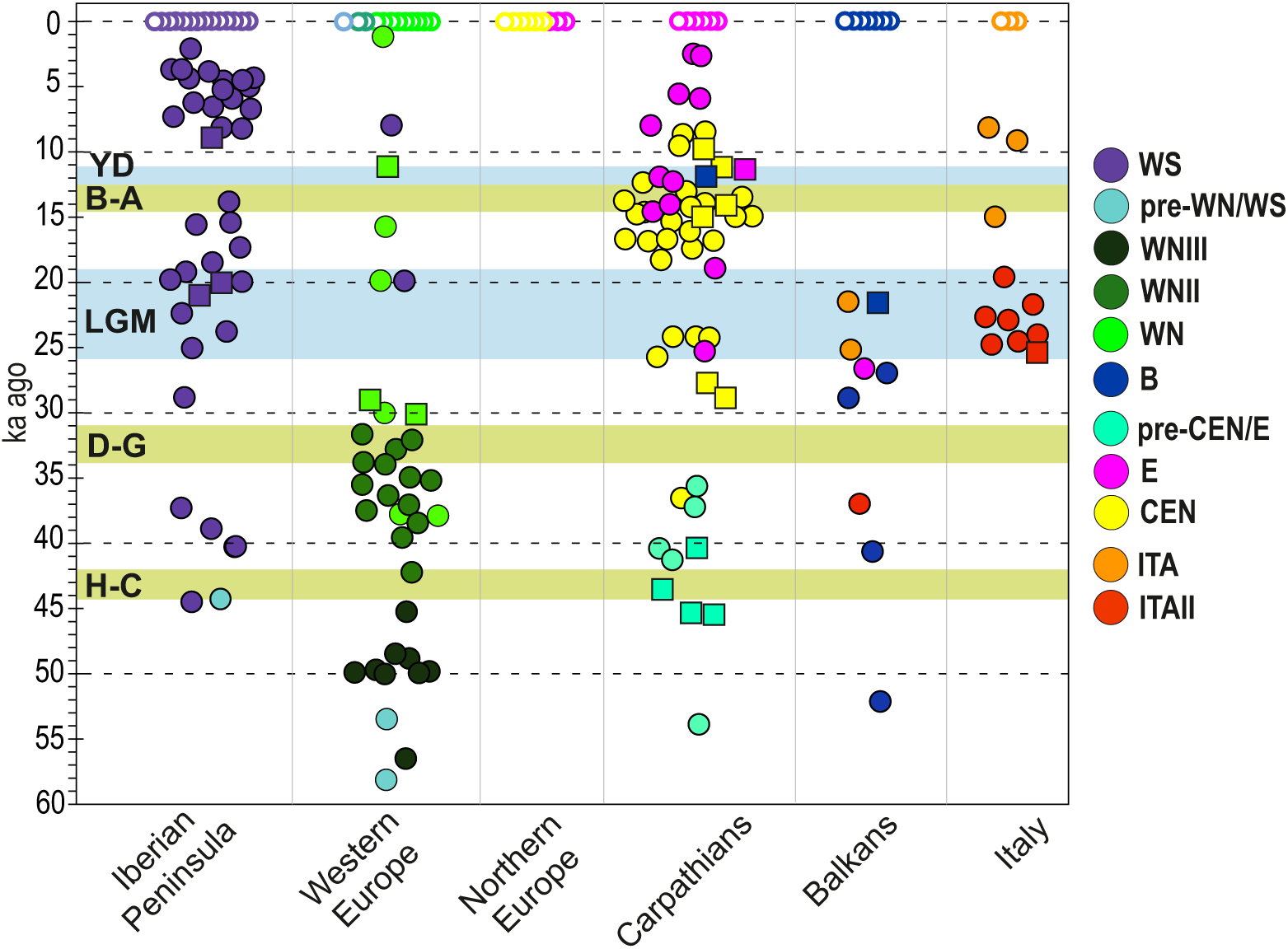
Temporal distribution of the mtDNA lineages divided by the geographical regions. Western Europe here includes France, southern Germany, Belgium, and the United Kingdom; the Northern Europe includes northern Germany and Poland (there were no ancient specimens from this region), the Carpathians include southern Poland, Czechia, Slovakia, Hungary, Ukraine and Romania and the Balkans include Bulgaria, Serbia and Croatia; Circles denote medians of age estimated using molecular approach while squares denote medians of calibrated radiocarbon ages. The green and blue strips indicate the main interstadials and stadials as in Figure 2.

The divergence of the main common vole mtDNA lineages estimated in this study was also older than previously suggested. All mtDNA lineages present in extant European populations (WS, WN, ITA, B, CEN, E) diverged prior to 40 ka ago, and the earliest specimens belonging to each of those lineages in our dataset pre-date 25 ka ago. This suggests, contrary to previous hypotheses (García et al., 2020; Heckel et al., 2005; Stojak et al., 2015), that climate deterioration during the LGM did not play the major role in the initial divergence of the main extant mtDNA lineages. However, the population decline and increased isolation during the period of the most inhospitable climatic conditions may have reinforced the previously existing divergence.

Most of the observed divergence events occurred between ca. 60 and 45 ka ago. They may be related to a long interstadial period identified in the palynological records across Europe: the Moershoofd Interstadial Complex in the Netherlands, the Pile Interstadial Complex at La Grande Pile pollen sequence (eastern France) and the Oerel and Glinde Interstadials at the Oerel pollen sequence (Northern Germany) dated to ca. 58–48 ka uncal BP and is usually correlated with Greenland Interstadials (GI) 16 and 14 (ca. 58–56.5 and 54–49.5 ka ago) (Helmens et al., 2014).

### 6.2 Phylogeography and demographic history of the common vole

#### 6.2.1 Western Europe and Iberian Peninsula

In Western Europe, we documented two consecutive mtDNA lineage turnovers at the end of MIS 3 (WNIII/WNII and WNII/WN). The first one took place around 45 ka ago (Figure 3). In our dataset there are few specimens of similar age from other parts of Europe; however, the divergent position of three specimens from the Western Carpathians (MI299, MI300, and MI1701; Figure 2) dated to 53 and 45 ka ago suggests that synchronous lineage turnovers may also have taken place in other parts of Europe. The second one occurred around 32 ka ago and appears to have been restricted to Western Europe (Figure 3). The oxygen isotope record from Greenland ice cores shows that the period between 45 and 29 ka ago was characterised by multiple short-term climatic oscillations (Rasmussen et al., 2014). Palynological records revealed two main interstadials which stand out during this period and have been identified in most of sediment sequences across Europe (Helmens, 2014). The older one, named Hengelo from its type locality in the Netherlands or Charbon in the Grande Pile sequence from eastern France, took place around 43–41 ka cal BP (38–36 ka uncal BP) (Helmens, 2014; Vandenberghe & van der Plicht, 2016), whilst the younger one (Denekamp – Grand Bois) occurred around 34–33 ka cal BP (31–29 ka uncal BP) (Guiter et al., 2003; Helmens, 2014), approximately at the same time as the recorded mtDNA lineage turnovers. The exact correlation of palynological data with Greenland ice-core records is problematic due to potential offsets and the wide error ranges involved. Hengelo–Charbon is usually associated with GI-11 (ca. 43.3–42.2 ka ago) or GI-10 (ca. 41.5–40.8 ka ago), although at times to the earlier GI-12 (ca. 46.8–44.2 ka ago) as the longest and most pronounced interstadial around the end of MIS 3, and Denekamp–Grand Bois with GI-8 (38.2–36.6 ka ago) (Helmens, 2014). Both interstadials were characterised by the emergence of *Betula* and *Pinus* forests in Western Europe, and of *Betula, Larix* and, *Pinus* forests in Central Europe, although it is assumed that the landscape remained relatively open as the duration of these interstadials was too short for the development of full forest cover (Guiter et al., 2003; Helmens, 2014). It was shown, however, that even partitioned landscapes limit dispersal and promote local extinctions of common vole populations (Delattre, Giraudoux, Baudry, Quere, & Fichet-Calvet, 1996), thus fragmentation of primarily open, stadial habitats into patchy and mosaic interstadial landscapes might have led to large scale decreases of population density and local or regional extinctions. A similar explanation for common vole population dynamics was previously suggested by Tougard et al. (2008). Martínková et al. (2013) showed partial replacement of mtDNA within the WN lineage in Late Holocene common vole populations from northern France and Belgium, and also suggested that the main factor driving this process was likely landscape reorganization.

The record from Northern Spain, occupied by the WS lineage, suggests population continuity throughout the last ca. 45 ka. The demographic reconstruction showed a drastic reduction of the effective population size around the Pleistocene/Holocene transition (Figures S2 and S3). This is in agreement with previous findings based on mtDNA cytochrome *b* and a smaller sample size (Baca et al., 2020). Palynological records from the region from which both modern and ancient specimens originated suggest that during the Bølling–Allerød interstadial (14.7–12.8 ka ago) a high proportion of open landscapes persisted until the expansion of deciduous woodlands started in the Early Holocene (Carrión et al., 2010).

#### 6.2.2 Central and South-eastern Europe

The survival of common vole populations throughout the LGM at high latitudes, including the Carpathians, has been previously suggested based on multiple lines of evidence. The fossil record suggests the continuous presence of the common vole in the Pannonian Basin (Pazonyi, 2004) and even north of the Carpathians (Sommer & Nadachowski, 2006), although these findings were based only on the stratigraphic position of the specimens, rather than direct dates. The Carpathians was also suggested as a northern refugium based on the distribution of the Eastern mtDNA haplogroup in modern populations (Stojak et al., 2016, 2015) and ecological niche modelling (Stojak, Borowik, Górny, McDevitt, & Wójcik, 2019). Our data support a northern survival of the common vole throughout the LGM. In the Western Carpathians, the northernmost part of this mountain range, we found all individuals with ages estimated, or directly dated, to between 36 and 10 ka ago to belong to the CEN mtDNA lineage. Two specimens from Šarkanica yielded pre-LGM, direct radiocarbon dates (28.9 and 27.8 ka cal BP; Table S3) while the age of a further four specimens was estimated to between 25.9 and 24.2 ka ago (Figure 2, Table S1). Recently, Lemanik et al. (2020) found a signal of rapid growth of the effective female population size (Nef) of the common vole population from the Western Carpathians starting ca. 21 ka cal BP that continued until ca. 15 ka cal BP. Together, these results are consistent with population continuity through the LGM, although accompanied by a significant reduction of population size.

Most of Central, Eastern, and Southeastern Europe is at present occupied by the Eastern lineage. It was suggested, based on both modern and ancient DNA, that the expansion of this lineage started around the Pleistocene/Holocene transition (Baca et al., 2020; Stojak et al., 2016). Our new data allow us to refine the history of this lineage. Prior to the maximal extension of the Scandinavian Ice Sheet (23–19 ka ago), we detected the Eastern lineage only in Southwestern Romania and Bulgaria. The presence of younger specimens bearing this mtDNA lineage in the same area, dated to 18.8 and 14 ka ago, suggests that this lineage may have survived the LGM in this part of the Carpathians. The expansion of the Eastern lineage from the South-eastern Carpathian area is also consistent with the finding of the highest genetic diversity in the extant common vole populations in this area (Stojak et al., 2016). The colonization of vast territories of Central and Eastern Europe, and replacement of Central and Balkan mtDNA lineages, took place somewhat later during the Bølling–Allerød or the Younger Dryas. In contrast to previous suggestions (Baca et al., 2020), the similar ages of Central and Eastern lineage individuals from the Muráň 3 and Býčí sites estimated here, suggest that both lineages may have coexisted in the area for some time (Figure 2, Table S1), prior to the final extirpation of the Central lineage in the Early Holocene. Although, the limited resolution of molecular age estimation and lack of possibility to track admixture with mtDNA does not allow for fine scale reconstruction of this process.

The record from the Italian peninsula suggests a replacement of the ITAII lineage with the ITA lineage which survived in Northern Italy and Switzerland until the present day. Although the available data come from sites with a limited temporal span (Appendix A), our dated phylogeny suggests that the extirpation of the ITAII lineage took place at some point after the LGM and the expansion of the ITA lineage occurred no later than the Bølling–Allerød warming. This lineage turnover may be reflected in the significant decrease in common vole remains for a short period after the LGM which was observed in the southern Italian peninsula (Berto, López-García, & Luzi, 2019). This is more consistent with repeated southward expansions of subsequent common vole populations rather than with their continuous presence in the region.

#### 6.2.3 Comparison with other European Late Pleistocene species

The common vole, along with the collared lemming and narrow-headed voles, are amongst the most numerous small mammals found in the assemblages from the last glacial period in Europe and are assumed to have coexisted during much of this period. The explanation for this paradox, in which a temperate species coexisted with cold-adapted ones, is the high tolerance of common vole to low temperatures (Tougard et al., 2008). Our study corroborates the continuous presence of the species at middle and high latitudes of Europe throughout the last at least 60 ka (Figure 3), although the evolutionary history of the common vole differed in some respects from other species inhabiting Europe during the Late Pleistocene. The long-term regional continuity of the main lineages of common vole with limited evidence for migrations suggested by our mtDNA analysis is in contrast with the Late Pleistocene evolutionary histories of megafaunal species such as cave bears and mammoths, both of which showed evidence for long-distance migrations and large-scale population replacements (Fellows Yates et al., 2017; Gretzinger et al., 2019). The collared lemming also showed a distinct pattern of mtDNA diversity consistent with multiple continent-wide population replacements (Palkopoulou et al., 2016). This was likely related to differences in mobility of the two species, with collared lemmings being more capable of long-distance dispersals than the common vole (Ehrich et al., 2001). On the other hand, available data suggest that despite this apparently different, species-specific history, common vole populations were affected by the same climatic and environmental changes as the cold adapted taxa. The extirpation of the WNIII lineage in Western Europe, and potentially of other common vole populations across Europe, took place around the same time as the continent-wide disappearance of collared lemming populations represented by their mtDNA lineages 1 and 2 (Palkopoulou et al., 2016). Similarly, the disappearance of the WNII lineage of the common voles around 32 ka ago is at approximately the same time as the population replacement of cave bears recorded in the Ach Valley, Germany (Münzel et al., 2011), and of woolly mammoth populations across the whole of Europe (Fellows Yates et al., 2017; Palkopoulou et al., 2013). Around 32 ka ago a new population of collared lemming (mtDNA lineage 3) appeared in Europe after a potential short-term extirpation (Palkopoulou et al., 2016). It was suggested that the main driver affecting mammalian species throughout the last glaciation were the abrupt warmings occurring at the onset of the Greenland Interstadials (Cooper et al., 2015). Nevertheless, whether it was the abruptness of the climatic changes, or the subsequent emergence of interstadial environments is unclear, although the high environmental instability in the period between 45 and 29 ka ago appears to have influenced a whole spectrum of differently adapted species including the common vole.

The other period that seems to have had a large impact on common vole populations was the Pleistocene/Holocene transition. The climatic warming and the emergence of forests during the Bølling–Allerød interstadial (14.7–12.8 ka ago) and in the Early Holocene (11.7–9 ka ago) are considered a main causes of extirpation of many cold-adapted species from Europe including the collared lemming and narrow-headed vole (Berto, Szymanek, et al., 2021; Royer et al., 2016). Our data confirmed previous findings indicating a substantial population decline of the common vole in Northern Spain, and mtDNA lineage replacement in the Western Carpathians (Baca et al., 2020). In addition, we identified a potential lineage turnover on the Italian peninsula. This, together with a probable extirpation of the common vole from the British Isles in the Early Holocene (Baca et al., 2020), suggests a continent-wide impact of interstadial environments on common vole populations.

## 7 Conclusions

Our study shows that the initial diversification of Last Glacial and extant common vole took place around 90 ka ago, during the Brørup interstadial (MIS 5c, GI-23). The divergence of mtDNA lineages present in extant common vole populations, as well as the first appearance of specimens belonging to those lineages, occurred earlier than previously estimated, mostly during the MIS 3 (57–29 ka ago). At high latitudes of Europe, we detected lineage turnovers whose dating suggest that they were likely caused by the fragmentation of primary habitats through reforestation during the interstadials occurring between 45 and 29 ka ago. In comparison, the climate deterioration during the LGM appears to have had a milder effect on common vole populations. More recent demographic changes and lineage turnovers like those recorded in Spain, the Western Carpathians, and Italy took place after the LGM or around the Pleistocene/Holocene transition and were also likely related to climatic warming. Altogether, this suggests that during the last glacial period, the evolutionary history of the common vole was distinct from typical cold-adapted species associated with steppe-tundra environments, although they responded to similar climatic and environmental changes. However, in contrast to the collared lemming and narrow-headed vole, the common vole did not become extinct in Europe at the end of the Pleistocene, possibly due to higher ecological plasticity, and eventually expanded during the mid-Holocene, taking advantage of secondary habitats such as agricultural fields. Overall, this suggests that habitat availability, rather than climatic variables, is the primary factor affecting common vole populations.

## 2 Acknowledgements

This research was supported by the Polish National Science Centre grants no.: 2015/19/D/NZ8/03878 to MB and 2017/25/B/NZ8/02005 to AN. Partial funding came from grant 31003A_176209 from the Swiss National Science Foundation to GH. XM was supported by the IT930-16 grant from Basque Science System (Basque Government). Fieldwork at Rocen-Pail (France) was granted by the French Ministry of Culture (MCC) through the Pays-de-la-Loire Regional Archaeology Service (DRAC/SRA) and in 2016 by the Mécène & Loire Fundation (http://www.mecene-et-loire.fr/). JML-G was supported by a Ramón y Cajal contract (RYC-2016-19386) with financial sponsorship from the Spanish Ministry of Science and Innovation. EL was supported by the Alexander von Humboldt Foundation with a Humboldt Research Fellowship for postdoctoral researchers (ESP1209403HFST-P) Analysis of modern Spanish specimens was supported by the Spanish Ministry of Science and Innovation and European Regional Development Fund, projects CGL2011-30274 and CGL2015-71255-P (MINECO-FEDER, EU). ST has received funding from the European Research Council under the European Union’s Horizon 2020 Research and Innovation Programme (grant agreement No. 803147 RESOLUTION, https://site.unibo.it/resolution-erc/en). AP acknowledges funding by the Romanian Research Authority (UEFISCDI) through grants PCCF 16/2016 (DARKFOOD), PCE 2282/2020 (ECHOES), and EEA Grant 126/2018 (KARSTHIVES). We also acknowledge the late Rebbeca Miller the director of Trou Al’Wesse excavations and the “AWaP – Agence Wallonne du Patrimoine” as the main funding institution of the work at the site

## 10 Data Availability

The consensus mtDNA sequences generated in this study have been deposited in GenBank under accession numbers OL588336 - OL588524. The alignment used for the reconstruction of phylogeny have been deposited in Dryad (doi:10.5061/dryad.4j0zpc8d9). Mitochondrial alignments generated in this study have been deposited in the European Nucleotide Archive under project number PRJEB53474.

## 11 Author Contributions

MB, DP and AN designed research; MB, DP, AL, HF and XW performed research; MB analysed the data and prepared figures; SBC, NC, GCB, ED, JTG, GH, IH, MVK, LL, JMLG, EL, ZM, JML, XM, AP, VP, TH, SER, BR, AR, JRS. JS, ST and JMW contributed samples; MB and AN wrote the paper with the input from all co-authors.

## References

Baca, M., Nadachowski, A., Lipecki, G., Mackiewicz, P., Marciszak, A., Popović, D., … Wojtal, P. (2017). Impact of climatic changes in the Late Pleistocene on migrations and extinction of mammals in Europe: four case studies. Geological Quarterly, 61(2), 291–304. doi: 10.7306/gq.1319

Baca, M., Popović, D., Baca, K., Lemanik, A., Doan, K., Horáček, I., … Nadachowski, A. (2020). Diverse responses of common vole (Microtus arvalis) populations to Late Glacial and Early Holocene climate changes – Evidence from ancient DNA. Quaternary Science Reviews, 233, 106239. doi: 10.1016/j.quascirev.2020.106239

Baca, M., Popović, D., Lemanik, A., Baca, K., Horáček, I., & Nadachowski, A. (2019). Highly divergent lineage of narrow-headed vole from the Late Pleistocene Europe. Scientific Reports, 9(1), 17799. doi: 10.1038/s41598-019-53937-1

Baca, M., Popović, D., Lemanik, A., Fewlass, H., Talamo, S., Zima, J., Ridush, B., Popov, V., & Nadachowski, A. (2021) The Tien Shan vole (Microtus ilaeus; Rodentia: Cricetidae) as a new species in the Late Pleistocene of Europe. Ecology and Evolution, 11, 16113–16125

Berto, C., López-García, J. M., & Luzi, E. (2019). Changes in the Late Pleistocene small-mammal distribution in the Italian Peninsula. Quaternary Science Reviews, 225, 106019. doi: 10.1016/j.quascirev.2019.106019

Berto, C., Nadachowski, A., Pereswiet-Soltan, A., Lemanik, A., & Kot, M. (2021). The Middle Pleistocene small mammals from the lower layers of Tunel Wielki Cave (Kraków-Częstochowa Upland): An Early Toringian assemblage in Poland. Quaternary International, 577, 52–70. doi: 10.1016/j.quaint.2020.10.023

Berto, C., Szymanek, M., Blain, H.-A., Pereswiet-Soltan, A., Krajcarz, M., & Kot, M. (in press). Small vertebrate and mollusc community response to the latest Pleistocene-Holocene environment and climate changes in the Kraków-Częstochowa Upland (Poland, Central Europe). Quaternary International, doi: 10.1016/j.quaint.2021.09.010

Beysard, M., & Heckel, G. (2014). Structure and dynamics of hybrid zones at different stages of speciation in the common vole (Microtus arvalis). Molecular Ecology, 23(3), 673–687. doi: 10.1111/mec.12613

Braaker, S., & Heckel, G. (2009). Transalpine colonisation and partial phylogeographic erosion by dispersal in the common vole (Microtus arvalis). Molecular Ecology, 18(11), 2528–2531. doi: 10.1111/j.1365-294X.2009.04189.x

Bronk Ramsey, C. (2009). Bayesian Analysis of Radiocarbon Dates. Radiocarbon, 51(1), 337–360. doi: 10.1017/S0033822200033865

Bužan, E. V., Förster, D. W., Searle, J. B., & Kryštufek, B. (2010). A new cytochrome b phylogroup of the common vole (Microtus arvalis) endemic to the Balkans and its implications for the evolutionary history of the species. Biological Journal of the Linnean Society, 100(4), 788–796. doi: 10.1111/j.1095-8312.2010.01451.x

Carrión, J. S., Fernández, S., González-Sampériz, P., López-Merino, L., Carrión-Marco, Y., Gil-Romera, G., … Burjachs, F. (2010). Expected trends and surprises in the Lateglacial and Holocene vegetation history of the Iberian Peninsula and Balearic Islands. Review of Palaeobotany and Palynology, 162(3), 458–475. doi: 10.1016/j.revpalbo.2009.12.007

Chaline, J. (1972). Les rongeures du pléistocène moyen et supérieur de France:(systématique, biostratigraphie, paléoclimatologie). In Cahiers Paléontologie. Paris: CNRS.

Cooper, A., Turney, C., Hughen, K. A., Barry, W., McDonald, H. G., & Bradshaw, C. J. A. (2015). Abrupt warming events drove Late Pleistocene Holarctic megafaunal turnover. Science, 349, 1–8. doi: 10.1126/science.aac4315

Dabney, J., Knapp, M., Glocke, I., Gansauge, M.-T., Weihmann, A., Nickel, B., … Meyer, M. (2013). Complete mitochondrial genome sequence of a Middle Pleistocene cave bear reconstructed from ultrashort DNA fragments. Proceedings of the National Academy of Sciences of the United States of America, 110(39), 15758–15763. doi: 10.1073/pnas.1314445110

Darriba, D., Taboada, G. L., Doallo, R., & Posada, D. (2012). jModelTest 2: more models, new heuristics and parallel computing. Nature Methods, 9(8), 772–772. doi: 10.1038/nmeth.2109

Delattre, P., Giraudoux, P., Baudry, J., Quere, J. P., & Fichet-Calvet, E. (1996). Effect of landscape structure on Common Vole (Microtus arvalis) distribution and abundance at several space scales. Landscape Ecology, 288(5), 279–288.

Duchene, S., Lemey, P., Stadler, T., Ho, S. Y. W., Duchene, D. A., Dhanasekaran, V., & Baele, G. (2020). Bayesian evaluation of temporal signal in measurably evolving populations. Molecular Biology and Evolution, 37(11), 3363–3379. doi: 10.1093/molbev/msaa163

Ehrich, D., Jorde, P. E., Krebs, C. J., Kenney, A. J., Stacy, J. E., & Stenseth, N. C. (2001). Spatial structure of lemming populations (Dicrostonyx groenlandicus) fluctuating in density. Molecular Ecology, 10(2), 481–495. doi: 10.1046/j.1365-294X.2001.01229.x

Fellows Yates, J. A., Drucker, D. G., Reiter, E., Heumos, S., Welker, F., Münzel, S. C., … Krause, J. (2017). Central European Woolly Mammoth population dynamics: insights from Late Pleistocene mitochondrial genomes. Scientific Reports, 7(1), 17714. doi: 10.1038/s41598-017-17723-1

Fewlass, H., Tuna, T., Fagault, Y., Hublin, J. J., Kromer, B., Bard, E., & Talamo, S. (2019). Pretreatment and gaseous radiocarbon dating of 40–100 mg archaeological bone. Scientific Reports, 9(1), 1–11. doi: 10.1038/s41598-019-41557-8

Fink, S., Excoffier, L., & Heckel, G. (2004). Mitochondrial gene diversity in the common vole Microtus arvalis shaped by historical divergence and local adaptations. Molecular Ecology, 13, 3501–3514. doi: 10.1111/j.1365-294X.2004.02351.x

Fischer, M. C., Foll, M., Heckel, G., & Excoffier, L. (2014). Continental-scale footprint of balancing and positive selection in a small rodent (Microtus arvalis). PLoS ONE, 9(11), e112332. doi: 10.1371/journal.pone.0112332

Folkertsma, R., Westbury, M. V., Eccard, J.A., & Hofreiter, M. (2018) The complete mitochondrial genome of the common vole, Microtus arvalis (Rodentia: Arvicolinae). Mitochondrial DNA Part B: Resources, 3, 446–447.

Gansauge, M. T., Aximu-Petri, A., Nagel, S., & Meyer, M. (2020). Manual and automated preparation of single-stranded DNA libraries for the sequencing of DNA from ancient biological remains and other sources of highly degraded DNA. Nature Protocols, 15(8), 2279–2300. doi: 10.1038/s41596-020-0338-0

García, J. T., Domínguez-Villaseñor, J., Alda, F., Calero-Riestra, M., Pérez Olea, P., Fargallo, J. A., … Viñuela, J. (2020). A complex scenario of glacial survival in Mediterranean and continental refugia of a temperate continental vole species (Microtus arvalis) in Europe. Journal of Zoological Systematics and Evolutionary Research, 58(1), 459–474. doi: 10.1111/jzs.12323

Gretzinger, J., Molak, M., Reiter, E., Pfrengle, S., Urban, C., Neukamm, J., … Schuenemann, V. J. (2019). Large-scale mitogenomic analysis of the phylogeography of the Late Pleistocene cave bear. Scientific Reports, 9(1), 1–11. doi: 10.1038/s41598-019-47073-z

Guiter, F., Andrieu-Ponel, V., de Beaulieu, J.-L., Cheddadi, R., Calvez, M., Ponel, P., … Goeury, C. (2003). The last climatic cycles in Western Europe : a comparison between long continuous lacustrine sequences from France and other terrestrial records. Quaternary International, 111, 59–74. doi: 10.1016/S1040-6182(03)00015-6

Haynes, S., Jaarola, M., & Searle, J. B. (2003). Phylogeography of the common vole (Microtus arvalis) with particular emphasis on the colonization of the Orkney archipelago. Molecular Ecology, 12(4), 951–956. doi: 10.1046/j.1365-294X.2003.01795.x

Heckel, G., Burri, R., Fink, S., Desmet, J.-F., & Excoffier, L. (2005). Genetic structure and colonization processes in European populations of the common vole, Microtus arvalis. Evolution; International Journal of Organic Evolution, 59(10), 2231–2242.. doi: 10.1554/05-255.1.

Helmens, K. F. (2014). The Last Interglacial-Glacial cycle (MIS 5-2) re-examined based on long proxy records from central and northern Europe. Quaternary Science Reviews, 86, 115–123. doi: 10.1016/j.quascirev.2013.12.012

Horáček, I., & Ložek, V. (1988). Palaeozoology and the Mid-European Quaternary Past: Scope of the Approach and Selected Results. Praha: Rozpravy ČSAV, Řada MPV.

Jacob, J., Manson, P., Barfknecht, R., & Fredricks, T. (2014). Common vole (Microtus arvalis) ecology and management: Implications for risk assessment of plant protection products. Pest Management Science, Vol. 70, pp. 869–878. John Wiley & Sons, Ltd. doi: 10.1002/ps.3695

Jánossy, D. (1986). Pleistocene vertebrate faunas of Hungary. In Pleistocene vertebrate faunas of Hungary. Amsterdam-Oxford-New York-Takyo: Elsevier. doi: 10.1016/0047-2484(89)90045-6

Jónsson, H., Ginolhac, A., Schubert, M., Johnson, P. L. F., & Orlando, L. (2013). MapDamage2.0: Fast approximate Bayesian estimates of ancient DNA damage parameters. Bioinformatics, 29(13), 1682–1684. doi: 10.1093/bioinformatics/btt193

Katoh, K., & Standley, D. M. (2013). MAFFT multiple sequence alignment software version 7: Improvements in performance and usability. Molecular Biology and Evolution, 30(4), 772–780. doi: 10.1093/molbev/mst010

Kučera, J., Suvova, Z., & Horáček, I. (2009). Early Middle Pleistocene glacial community of rodents (Rodentia): Stránská skála cave (Czech Republic). Lynx, (40), 43–69.

Lemanik, A., Baca, M., Wertz, K., Socha, P., Popović, D., Tomek, T., … Nadachowski, A. (2020). The impact of major warming at 14.7 ka on environmental changes and activity of Final Palaeolithic hunters at a local scale (Orawa-Nowy Targ Basin, Western Carpathians, Poland). Archaeological and Anthropological Sciences, 12(3). doi: 10.1007/s12520-020-01020-6

Li, H., & Durbin, R. (2010). Fast and accurate long-read alignment with Burrows-Wheeler transform. Bioinformatics (Oxford, England), 26(5), 589–595. doi: 10.1093/bioinformatics/btp698

Li, H., Handsaker, B., Wysoker, A., Fennell, T., Ruan, J., Homer, N., … Subgroup, 1000 Genome Project Data Processing. (2009). The Sequence Alignment/Map format and SAMtools. Bioinformatics, 25(16), 2078–2079. doi: 10.1093/bioinformatics/btp352

Lischer, H. E. L., Excoffier, L., & Heckel, G. (2014). Ignoring heterozygous sites biases phylogenomic estimates of divergence times: Implications for the evolutionary history of Microtus voles. Molecular Biology and Evolution, 31(4), 817–831. doi: 10.1093/molbev/mst271

Lorenzen, E. D., Nogués-Bravo, D., Orlando, L., Weinstock, J., Binladen, J., Marske, K. A., … Willerslev, E. (2011). Species-specific responses of Late Quaternary megafauna to climate and humans. Nature, 479(7373), 359–364. doi: 10.1038/nature10574

Martínková, N., Barnett, R., Cucchi, T., Struchen, R., Pascal, M., Pascal, M., … Searle, J. B. (2013). Divergent evolutionary processes associated with colonization of offshore islands. Molecular Ecology, 22(20), 5205–5220. doi: 10.1111/mec.12462

Maul, L. C., & Markova, A. K. (2007). Similarity and regional differences in Quaternary arvicolid evolution in Central and Eastern Europe. Quaternary International, 160(1), 81–99. doi: 10.1016/j.quaint.2006.09.010

Meyer, M., & Kircher, M. (2010). Illumina sequencing library preparation for highly multiplexed target capture and sequencing. Cold Spring Harbor Protocols, 5(6), t5448. doi: 10.1101/pdb.prot5448

Milne, I., Stephen, G., Bayer, M., Cock, P. J. A., Pritchard, L., Cardle, L., … Marshall, D. (2013). Using Tablet for visual exploration of second-generation sequencing data. Briefings in Bioinformatics, 14(2), 193–202. doi: 10.1093/bib/bbs012

Mourier, T., Ho, S.Y.W., Gilbert, M.T.P., Willerslev, E., & Orlando, L. (2012) Statistical guidelines for detecting past population shifts using ancient DNA. Molecular Biology and Evolution, 29, 2241–51. doi:10.1093/molbev/mss094

Münzel, S. C., Stiller, M., Hofreiter, M., Mittnik, A., Conard, N. J., & Bocherens, H. (2011). Pleistocene bears in the Swabian Jura (Germany): Genetic replacement, ecological displacement, extinctions and survival. Quaternary International, 245(2), 1–13. doi: 10.1016/j.quaint.2011.03.060

Nadachowski, A. (1989). Origin and history of the present rodent fauna in Poland based on fossil evidence. Acta Theriologica, 34(2), 37–53.

Palkopoulou, E., Baca, M., Abramson, N. I., Sablin, M., Socha, P., Nadachowski, A., … Dalén, L. (2016). Synchronous genetic turnovers across Western Eurasia in Late Pleistocene collared lemmings. Global Change Biology, 22(5), 1710–1721. doi: 10.1111/gcb.13214

Palkopoulou, E., Dalen, L., Lister, A. M., Vartanyan, S., Sablin, M., Sher, A., … Thomas, J. A. (2013). Holarctic genetic structure and range dynamics in the woolly mammoth. Proceedings of the Royal Society B: Biological Sciences, 280, 20131910. doi: 10.1098/rspb.2013.1910.

Pazonyi, P. (2004). Mammalian ecosystem dynamics in the Carpathian Basin during the last 27,000 years. Palaeogeography, Palaeoclimatology, Palaeoecology, 212(3–4), 295–314. doi: 10.1016/j.palaeo.2004.06.008

Rasmussen, S. O., Bigler, M., Blockley, S. P., Blunier, T., Buchardt, S. L., Clausen, H. B., … Winstrup, M. (2014). A stratigraphic framework for abrupt climatic changes during the Last Glacial period based on three synchronized Greenland ice-core records: refining and extending the INTIMATE event stratigraphy. Quaternary Science Reviews, 106, 14–28. doi: 10.1016/j.quascirev.2014.09.007

Reimer, P. J., Austin, W. E. N., Bard, E., Bayliss, A., Blackwell, P. G., Bronk Ramsey, C., … Talamo, S. (2020). The IntCal20 northern hemisphere radiocarbon age calibration curve (0–55 cal kBP). Radiocarbon, 62(4), 725–757. doi: 10.1017/rdc.2020.41

Richard, M., Falguères, C., Valladas, H., Ghaleb, B., Pons-Branchu, E., Mercier, N., … Conard, N. J. (2019). New electron spin resonance (ESR) ages from Geißenklösterle Cave: A chronological study of the Middle and early Upper Paleolithic layers. Journal of Human Evolution, 133, 133–145. doi: 10.1016/j.jhevol.2019.05.014

Royer, A., Montuire, S., Legendre, S., Discamps, E., Jeannet, M., & Lécuyer, C. (2016). Investigating the influence of climate changes on rodent communities at a regional-scale (MIS 1-3, Southwestern France). PLoS ONE, 11(1), e0145600. doi: 10.1371/journal.pone.0145600

Schubert, M., Lindgreen, S., & Orlando, L. (2016). AdapterRemoval v2: Rapid adapter trimming, identification, and read merging. BMC Research Notes, 9(1), 1–7. doi: 10.1186/s13104-016-1900-2

Socha, P. (2014). Rodent palaeofaunas from Biśnik Cave (Kraków-Częstochowa Upland, Poland): Palaeoecological, palaeoclimatic and biostratigraphic reconstruction. Quaternary International, 326–327, 64–81. doi: 10.1016/j.quaint.2013.12.027

Sommer, R. S., & Nadachowski, A. (2006). Glacial refugia of mammals in Europe: evidence from fossil records. Mammal Review, 36(4), 251–265. doi: 10.1111/j.1365-2907.2006.00093.x

Stewart, J. R., Lister, A. M., Barnes, I., & Dalén, L. (2010). Refugia revisited: individualistic responses of species in space and time. Proceedings. Biological Sciences / The Royal Society, 277(1682), 661–671. doi: 10.1098/rspb.2009.1272

Stojak, J., Borowik, T., Górny, M., McDevitt, A. D., & Wójcik, J. M. (2019). Climatic influences on the genetic structure and distribution of the common vole and field vole in Europe. Mammal Research, 64(1), 19–29. doi: 10.1007/s13364-018-0395-8

Stojak, J., McDevitt, A. D., Herman, J. S., Kryštufek, B., Uhlíková, J., Purger, J. J., … Wójcik, J. M. (2016). Between the Balkans and the Baltic: Phylogeography of a Common Vole mitochondrial DNA lineage limited to Central Europe. PLOS ONE, 11(12), e0168621. doi: 10.1371/journal.pone.0168621

Stojak, J., Mcdevitt, A. D., Herman, J. S., Searle, J. B., & Wójcik, J. M. (2015). Post-glacial colonization of eastern Europe from the Carpathian refugium: evidence from mitochondrial DNA of the common vole Microtus arvalis. Biological Journal of the Linnean Society, 115(4), 927–939.

Suchard, M. A., Lemey, P., Baele, G., Ayres, D. L., Drummond, A. J., & Rambaut, A. (2018). Bayesian phylogenetic and phylodynamic data integration using BEAST 1.10. Virus Evolution, 4(1), 1–5. doi: 10.1093/ve/vey016

Tougard, C., Renvoisé, E., Petitjean, A., & Quéré, J.-P. (2008). New insight into the colonization processes of common voles: inferences from molecular and fossil evidence. PloS One, 3(10), e3532. doi: 10.1371/journal.pone.0003532

Van Klinken, G. J. (1999). Bone collagen quality indicators for palaeodietary and radiocarbon measurements. Journal of Archaeological Science, 26(6), 687–695. doi: 10.1006/jasc.1998.0385

Vandenberghe, J., & van der Plicht, J. (2016). The age of the Hengelo interstadial revisited. Quaternary Geochronology, 32, 21–28. doi: 10.1016/j.quageo.2015.12.004

Wacker, L., Bonani, G., Friedrich, M., Hajdas, I., Kromer, B., Němec, M., … Vockenhuber, C. (2010). MICADAS: Routine and high-precision radiocarbon dating. Radiocarbon, 52(2), 252–262. doi: 10.1017/S0033822200045288

Wacker, L., Fahrni, S. M., Hajdas, I., Molnar, M., Synal, H. A., Szidat, S., & Zhang, Y. L. (2013). A versatile gas interface for routine radiocarbon analysis with a gas ion source. Nuclear Instruments and Methods in Physics Research, Section B: Beam Interactions with Materials and Atoms, 294, 315–319. doi: 10.1016/j.nimb.2012.02.009

Wacker, L., Němec, M., & Bourquin, J. (2010). A revolutionary graphitisation system: Fully automated, compact and simple. Nuclear Instruments and Methods in Physics Research, Section B: Beam Interactions with Materials and Atoms, 268(7–8), 931–934. doi: 10.1016/j.nimb.2009.10.067

Walther, G. R. (2010). Community and ecosystem responses to recent climate change. Philosophical Transactions of the Royal Society B: Biological Sciences, 365(1549), 2019–2024. doi: 10.1098/rstb.2010.0021

