## Supplementary Text S1 for "Ancient DNA reveals interstadials as a driver of the common vole population dynamics during the last glacial period"

^12^ Histoire Naturelle de l’Homme Préhistorique (HNHP), UMR 7194, Dept. Homme et Environnement du Muséum national d’Histoire Naturelle, MNHN-CNRS-UPVD, Musée de l’Homme, Paris, France

^13^ Institut Català de Paleoecologia Humana i Evolució Social (IPHES-CERCA), Tarragona, Spain

^14^ Natural History Museum, Belgrade, Serbia

^15^ Institute for Quaternary Palaeontology and Geology, Croatian Academy of Sciences and Arts, Zagreb, Croatia

^16^ Department of Geology University of the Basque Country UPV/EHU, Bilbao, Spain

^17^ Service de Préhistoire, Université de Liège, Place du 20 Août 7, 4000 Liège, Belgium

^18^ Institute of Speleology “E. Racovitza”, Bucharest, Romania & Romanian Institute of Science and Technology, Cluj-Napoca, Romania

^19^ Institute of Biodiversity and Ecosystem Research, Bulgarian Academy of Sciences, Sophia, Bulgaria

^20^ Interdisciplinary Center for Archaeology and Evolution of Human Behavior (ICArEHB), Universidade do Algarve, Faro, Portugal.

^21^ Department of Physical Geography, Geomorphology and Paleogeography, Yuriy Fed'kovych Chernivtsi National University, Ukraine

^22^ Biogéosciences*,* UMR 6282*,* CNRS*,* Université Bourgogne Franche*-*Comté*,* Dijon, France

^23^ Faculty of Science and Technology, Bournemouth University, Poole, United Kingdom

^24^ Mammal Research Institute**,** Polish Academy of Sciences, Białowieża, Poland

^25^ Department of Chemistry G. Ciamician, Alma Mater Studiorum, University of Bologna, Bologna, Italy

^+^ - these authors contributed equally

* Corresponding authors:

**List of Supplementary Figures**

Figure S1. Results of LOO analysis……………………………………………………………….8

Figure S2. Changes of effective female population size (Nef) of common vole……...…………10

Figure S3. Changes of effective female population size (Nef) of common vole………………...11

**List of Supplementary Tables**

Table S1. Information on ancient samples analysed in this study (see separate Excel file)

Table S2. Information on modern samples analysed in this study (see separate Excel file)

Table S3. Details of radiocarbon dating of vole mandibles (see separate Excel file)

Table S4. Primers used to generate mitogenomes from various vole species…………………… 5

Table S5. Blocking oligos used in hybridization reaction………………………………………...6

Table S6. Results of Bayesian evaluation of temporal signal……………………………………. 7

Table S7. Comparison of four alternative models tested in BETS analysis……………………....7

Table S8. Testing tree priors for ‘dated’ dataset using GSS………………………………………8

Table S9. Model testing for ‘joint’ dataset using GSS.……………………...…………………..10

Table S10. Radiocarbon dates obtained to improve stratigraphic information of sites………….13

### 1. Supporting Methods and Results

#### 1.1. Modern specimens

The DNA of modern common vole specimens was extracted and processed in four laboratories, including the Mammal Research Institute of the Polish Academy of Sciences in Białowieża, Poland (MRI), the Laboratory of Paleogenetics and Conservation Genetics, the Centre of New Technologies at the University of Warsaw in Warsaw Poland (LPCG), the Institute of Ecology and Evolution, University of Bern, Switzerland (IEE) and the Instituto de Investigación en Recursos Cinegéticos, Ciudad Real, Spain (IREC). Information on which laboratories processed which specimens is provided in Table S2. Total genomic DNA was extracted from the MRI/LPCG specimens using the Syngen Tissue DNA Mini Kit at the MRI (Stojak et al., 2015) and sequencing libraries were generated, enriched, and sequenced at the LPCG. Total genomic DNA of the IREC/LPCG specimens was extracted using a standard AcNH4 protocol at the IREC (García et al., 2020) and further processed at the LPCG.

The genomic DNA of the LPCG specimens labelled ‘target’ in Table S2, was inspected by 1% agarose gel electrophoresis and sonicated to a mean length of ca. 200 bp using the Covaris S220 sonicator. The sheared DNA was transformed into double-indexed, double-stranded libraries as described in Section 1.3 except that the indexing polymerase chain reaction (PCR) was run for 10–12 cycles instead of 19 cycles. The libraries were pooled and enriched for mtDNA as described in Section 1.4, except that a single round of enrichment was performed, and post-hybridisation amplification was run for 10 cycles.

Specimen labelled ‘PCR’ in Table S2 was used to generate enrichment bait (Section 1.4). The amplified mitogenome was fragmented using the Covaris S220 sonicator and transformed into a sequencing library as described in Section 1.3 except that 10 cycles of indexing PCR were run.

Total genomic DNA from IEE ethanol preserved tissues was extracted using a phenol-chloroform procedure. The DNA was checked by 1% agarose gel electrophoresis, and the DNA concentration and quality were measured with the NanoDrop 2000 Spectrophotometer. The Illumina TruSeq DNA PCR-Free Library Prep Kit was used to generate the libraries for whole-genome resequencing. The sequencing was performed with the Illumina Hiseq 2000 or Novaseq 6000 using the next generation sequencing platform of the University of Bern.

#### 1.2. Ancient DNA extraction

DNA extraction and the pre-PCR library preparation steps were performed in the dedicated ancient DNA laboratory at the LPCG. All teeth were thoroughly cleaned with ultra-pure water in a 2 ml tube, crushed with a pipette tip and incubated overnight in 1 ml of extraction buffer (0.5 M EDTA pH = 8.0; 0.5% N-laurylsarcosine; 0.1 mg Proteinase K) at 38°C with agitation. After the incubation, one part of the extraction buffer was combined with 13 parts of binding buffer (5 M guanidine hydrochloride and 40% isopropanol) and eluted through a MinElute silica column (Qiagen, Hilden, Germany). The silica suspension was washed twice with 750 µl of PE buffer (80% ethanol, 10 mM Tris-HCl pH 7.5), dried and eluted twice with 30 µl of pre-warmed EB buffer (10 mM Tris-Cl, pH 8.5).

#### 1.3. Double-stranded library preparation

The double-indexed, double-stranded sequencing libraries were constructed following the protocol of Meyer and Kircher (2010) with minor modifications described in Baca et al. (2019) using 20 µl of DNA extract as input. The blunt-end repair was performed in a 30 µl reaction containing 1× buffer Tango, 15 U T4 polynucleotide kinase (Thermo), 3U T4 DNA polymerase, 100 µDNTPs, and 1 mM ATP. The reaction was incubated for 15 min at 25°C, followed by 5 min at 12°C and 20 min at 95°C to inactivate the enzymes. An adaptor ligation step was performed by adding 10 µl of the adapter ligation mix directly to the blunt-end repair reaction resulting in a final reaction volume of 40 µl containing 1× T4 DNA ligase buffer, 5% PEG-4000, 5 U T4 DNA ligase (Thermo Scientific) and 1 µM of the P5 and P7 adapters. The reaction was incubated for 30 min at 22°C and purified using magnetic beads. Adapter fill-in was performed by adding 20 µl of purified ligation product to 15 µl of the reaction master mix resulting in a 35 µl reaction containing 9.6 U of BST polymerase (New England Biolabs, Ipswich, MA, USA), 1× Thermopol buffer and 0.25 µM of the dNTPs. The reactions were incubated in a thermocycler for 20 min at 37°C, followed by heat inactivation at 80°C for 20 min. The DNA libraries were amplified in three replicates in 25 µl reaction volumes containing 10 µl of adapter-ligated DNA, 1× AmpliTaq Gold 360 Master Mix (Thermo Scientific) and 0.2 µM of the P5 and P7 indexing primers under the following conditions: 95°C for 12 min, 19 cycles of 95°C for 30 s, 60°C for 30 s, 72°C for 1 min and 72°C for 10 min. Each indexing primer contained a 7-bp long index. The amplification replicates were combined and purified using magnetic beads. The libraries were visualised by 2% agarose gel electrophoresis and quantified with the Denoxiv spectrophotometer.

#### 1.4. Single-stranded library preparation

We prepared single-stranded libraries for five specimens with poorly preserved DNA in which the enriched double-stranded libraries yielded very few target DNA molecules. Twenty microliters of DNA extract were combined with 9 µl of water and 1 µl of USER enzyme (New England Biolabs) and incubated for 1 h at 37°C. Further steps strictly followed the protocol outlined by Gansauge et al. (2020). The appropriate number of PCR cycles was determined with the qPCR using Illumina Library Quantification kit before indexing (KAPA). Indexing PCR was run in duplicate using AccuPrime™ *Pfx* DNA Polymerase (Thermo Scientific). Amplified libraries were combined, purified using magnetic beads and subjected to the target enrichment procedure (Section 1.5).

#### 1.5. Target enrichment of mtDNA

Target enrichment was performed to enrich the libraries with vole mtDNA. Hybridisation bait was produced using the modern DNA of the following vole species: common vole (*M. arvalis*), field vole (*M. agrestis*), root vole (*Alexandromys oeconomus*), narrow-headed vole (*Lasiopodomys gregalis*) and bank vole (*Clethrionomys glareolus*). Total genomic DNA was extracted from tissue fragments using the Syngen Tissue DNA Mini Kit at MRI. The bait was prepared and the enrichment procedure was performed at LPCG. The mitogenomes were amplified in four overlapping fragments (Table S4) using PrimeSTAR GXL DNA Polymerase (Takara Bio, Shiga, Japan).

Table S4. Primers used to generate the mitogenomes from various vole species

| Primer ID | Sequence | Product length |
| --- | --- | --- |
| MICMT01F | TGCAAGCATCCCATAAACAA | 3.8 kb |
| MICMT01R | ATGGGCCCGATAGCTTTATT |  |
| MICMT02F | CAAAATTCTCCGTGCTACCC | 4.4 kb |
| MICMT02R | TTGTGTGGTTGGGGTAAATG |  |
| MICMT03F | CGCCTCTTTCATTACCCCTA | 4.2 kb |
| MICMT03R | TCYCAGCCGATGAAGAGTTG |  |
| MICMT04F | ACCCHAACCTAAACCGATTC | 4.5 kb |
| MICMT04R | ATAAGGCCAGGACCAAACCT |  |

Each fragment was amplified separately. The amplification reaction was performed in 50 µl comprised of 20–50 ng of genomic DNA, 1× PrimeSTAR GXL Buffer, dNTPs (200 µM each), 0.2 µM primers and 2.5 U PrimeSTAR GXL DNA Polymerase. The PCR conditions were 30 cycles at 98°C for 10 s, 55°C for 15 s and 68°C for 30 to 50 s depending on the target length.

The PCR products were mixed in equimolar ratios and sonicated to a length of ca. 200 bp using the Covaris S220 sonicator. The sonicated DNA from various species was pooled and converted into DNA bait following the protocol reported by Maricić et al. (2010). Target enrichment was performed in solution following the protocol of Horn (2012). Hybridisation was performed using the Oligo aCGH/ChiP-on-Chip Hybridisation Kit (Agilent Technologies, Palo Alto, CA, USA). Each reaction (50 µl) consisted of 12–15 µl pooled libraries (up to five libraries), 25 µl of 2× hybridisation buffer, 5 µl of blocking agent, 4 µl of blocking oligos (25 µM each, Table S5) and 1–3 µl of DNA bait. The quantities of the pooled libraries and DNA bait were adjusted so the library to bait ratio was 10 to 1. Hybridisation was carried out for 20–24 h at 65°C in a thermocycler. After the incubation, the hybridisation reaction was incubated for 20 min with 5 µl of streptavidin-coated beads (Dynabeads MyOne C1, Thermo Fisher) to immobilise the enriched libraries. The bead-immobilised libraries were washed five times with BWT buffer (see Horn 2012 for buffer composition), incubated for 2 min at 50°C with HWE buffer, washed once with BWT buffer, transferred to a new tube, washed once with TET and resuspended in 35 µl of TE buffer. To separate the enriched library from the bait, the mixture was incubated for 5 min at 95°C, the beads were collected on a magnet and the eluate was transferred to a new tube. The enriched library was amplified 15 cycles in three separate reactions of 20 µl each using Herculase II Fusion DNA polymerase (Agilent Technologies), which was purified using magnetic beads and subjected to the second round of hybridisation and amplification. Libraries from the ancient and modern specimens were never pooled in one hybridisation reaction. Multiple enriched library pools, either ancient or modern, were combined for sequencing ensuring that all P5 and P7 indices were unique in the pool, quantified using qPCR (Illumina Library Quantification kit, KAPA) and sequenced on the NextSeq550 at the CeNT UW NGS Core Facility using the 150 bp Mid Output kit and a 2× 75 bp sequencing scheme. A custom Read1 primer was used for the single-stranded libraries, as described by Gansauge et al. (2020).

Table S5. Blocking oligos used in the hybridisation reaction

| BO3.P7.part1.F | AGATCGGAAGAGCACACGTCTGAACTCCAGTCAC-phosphate |
| --- | --- |
| BO4.P7.part1.R | GTGACTGGAGTTCAGACGTGTGCTCTTCCGATCT-phosphate |
| BO5.P7.part2.F | ATCTCGTATGCCGTCTTCTGCTTG-phosphate |
| BO6.P7.part2.R | CAAGCAGAAGACGGCATACGAGAT-phosphate |
| BO7.P5.part1.F | AATGATACGGCGACCACCGAGATCTACAC-phosphate |
| BO8.P5.part1.R | GTGTAGATCTCGGTGGTCGCCGTATCATT-phosphate |
| BO09.P5.part2.F | ACACTCTTTCCCTACACGACGCTCTTCCGATCT-phosphate |
| BO10.P5.part2.R | AGATCGGAAGAGCGTCGTGTAGGGAAAGAGTGT-phosphate |

#### 1.6. Sequence processing

The raw reads were demultiplexed using bcl2fastq v. 2.19 (Illumina). Overlapping reads were collapsed and adaptor and quality trimmed (-trimns, -trimqualities) using AdapterRemoval v. 2.2.2 (Schubert, Lindgreen, & Orlando, 2016) with the following parameters: --collapse, --minalignmentlength 4, --trims, --trimqualities, --gzip, --basename ‘sample’. We indicated the sequence of the second adapter the single-stranded libraries as --adapter2 GGAAGAGCGTCGTGTAGGGAAAGAGTGT. The reads were mapped to a common vole reference using the BWA-MEM algorithm. We used either the common vole mtDNA reference sequence (NC_038176) or to a mtDNA sequence based on one of the extant specimens (WM176). Duplicated, short (<30 bp) and low mapping quality reads (mapq <30) were removed by *samtools* v.1.9 using the *samtools view-q 30*; *samtools sort* and *samtools rmdup* commands. Variants and consensus sequences were called using *bcftools* v.1.9 (Li et al., 2009) with the *mpileup* and *call* commands. BED file, which is used for masking low coverage positions, was generated using the *genomecov* command from BEDtools v. 2.27 (Quinlan & Hall, 2010) and filtered to retain only positions with coverage less than 3 using the *awk* script. Read alignments and vcf files were inspected manually using Tablet v. 1.17 software (Milne et al., 2013).

#### 1.7. Details of the phylogenetic analyses

The sequences used in this study were aligned in MAFFT (Katoh & Standley, 2013). The alignment was visually inspected, the variations in length around nucleotide homopolymers were trimmed and singleton indels were deleted. BEAST 1.10.4 software was used to reconstruct the phylogenetic relationships, estimate the divergence times of the common vole lineages and age the not directly dated specimens.

The Bayesian evaluation of temporal signal (BETS, Duchene et al., 2020) was utilised to check if the temporal signal within our dataset were sufficient to calibrate the molecular clock. In this analysis, 71 sequences, 51 from extant and 20 from directly radiocarbon-dated specimens, were used. The medians of the calibrated radiocarbon ages were set as the sequence sampling times. The support for the four models was compared. In the first two models, real sampling times were assigned to the sequences (heterochronous analysis), and then either a strict clock or an uncorrelated relaxed log-normal clock (UCLN) was used. In the two other models, the same sampling time (i.e. isochronous analysis) was used for all sequences, and either a strict clock or the UCLN clock was applied. A constant population size tree prior to all analyses and a CTMC rate reference prior for the heterochronous datasets were applied (Ferreira & Suchard, 2008). Each analysis was run 50 million steps sampled every 5,000 steps and with the first 5 million steps discarded as burn-in. Convergence and stationarity were inspected in Tracer 1.7 (ESS >200 for all parameters). The log marginal likelihood (MLE) of each model was estimated using the generalised stepping-stone (GSS) sampling approach (Baele et al., 2016). The MLE calculation was comprised of 50 path steps, each run for one million iterations. Two replicates of each BEAST analysis were performed. The BETS analysis strongly supported the model with the correct sampling times and the strict clock over the other models (2lnBF >9), suggesting that our dataset was suitable for calibrating the molecular clock (Tables S6 and S7).

Table S6. Bayesian evaluation of the temporal signal

| ID | Description | Replicate | logML | mean logML |
| --- | --- | --- | --- | --- |
| **Model I** | **Tips; strict clock, CTMC rate prior** | Rep I | −11956,5 | **−11956,41** |
|  |  | Rep II | −11956,33 |  |
| Model II | Tips; UCLN clock, CTMC rate prior | Rep I | −11959,99 | −11961,13 |
|  |  | Rep II | −11962,27 |  |
| Model III | No tips; strict clock | Rep I | −12069,85 | −12068,99 |
|  |  | Rep II | −12068,12 |  |
| Model IV | No tips; UCLN clock | Rep I | −12012,92 | −12014,28 |
|  |  | Rep II | −12015,64 |  |

Table S7. Comparison of the four alternative models tested in the BETS analysis

| Model comparison | 2lnBF |
| --- | --- |
| Model I vs. Model II | **9,436** |
| Model I vs. Model III | **225,144** |
| Model I vs. Model IV | **115,723** |

Next, we tested, which tree prior fit best to our dated dataset. A constant population size and Skygrid tree priors were compared using GSS MLE. The analyses were conducted the same as for the BETS analysis, and ‘positive’ support was found for the Skygrid tree prior (Table S8).

Table S8. Testing models for the ‘dated’ dataset using GSS

| Analysis details | logML (replicates) | Mean logML | 2lnBF |
| --- | --- | --- | --- |
| Dated dataset, strict clock, Skygrid tree prior, CTMC rate prior | −11954,362 | **−11953,937** | **4,95** |
|  | −11953,514 |  |  |
| Dated dataset, strict clock, constant pop. Size tree prior, CTMC rate prior | −11956,546 | −11956,177 |  |
|  | −11955,809 |  |  |

Next, a leave-one-out analysis was performed on the directly dated specimens to check the accuracy of the age estimates produced using the available calibration dataset. In this analysis, the age of each directly radiocarbon-dated specimen was estimated using all of the remaining directly dated and modern specimens to calibrate the molecular clock. We set a gamma prior (shape = 2; scale = 50,000) on the age of the directly dated specimen whose age was to be estimated using the molecular dating approach and increased the operator weight of the age estimate to 5. We set a Skygrid tree prior and used a CTMC rate reference prior. Each analysis was run for 50 million generations sampled every 5,000 steps and with the first 5 million generations discarded as burn-in. Convergence and stationarity were inspected in Tracer 1.7 (ESS >200 for all parameters).

In the leave-one-out analysis, the 95% HPD intervals of the estimated ages for most of the specimens overlapped with their respective radiocarbon ages (2-sigma ranges; Figure S1). The estimated ages of three specimens did not overlap with the calibrated radiocarbon ages. In the case of MI1337 from layer 4 of Muráň 3 site (Slovakia), the estimated age was older than the calibrated radiocarbon age; in the cases of MI074 from layer IIIa from Obłazowa (WE), Poland and MI1355 from Šarkanica, Czechia, the estimated ages were younger than the respective calibrated radiocarbon ages (Figure S1). In the case of specimens from Muráň 3 and Obłazowa (WE), the calibrated radiocarbon ages of the specimens were in good agreement with the stratigraphic positions of the specimens. There are five radiocarbon dates available from Šarkanica, ranging between 26 and 20 cal ka BP, and both medians of the estimated age (16.7 ka) and calibrated radiocarbon age of the specimen (28.4 cal ka BP) were outside this range (see Section 2.10). Specimens MI074 and MI1337 yielded high sequence coverages (>95% of the analysed fragment covered). MI1355 had lower coverage (82% of the analysed fragment covered) than the other specimens, which may have affected the age estimate. The authors of the tip-dating approach recognise that possible errors may have several sources and that they are often extremely difficult to distinguish (Shapiro et al., 2011).

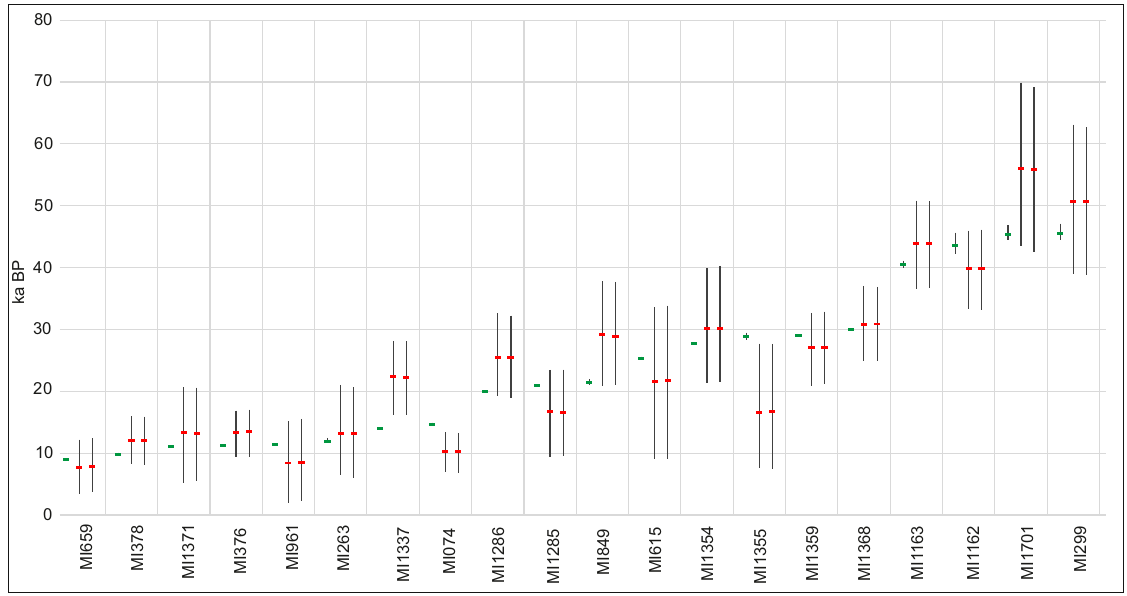

Figure S1. Results of the leave-one-out analysis. Three age estimates are presented for each specimen. The first one is a radiocarbon age (green dot denotes median calibrated age, and whiskers denote 94% probability range), whereas the next two are the two replicates of the molecular age estimates from BEAST (red dot denotes median age, and whiskers denote 95% HPD interval).

The results of the BETS and leave-one-out analyses suggested that our ‘dated’ dataset contained a sufficient phylogenetic signal to calibrate the molecular clock and accurately estimate the age of the not directly dated specimens. To estimate the age of each not directly dated specimen (n-128), we ran a separate BEAST analysis using the sequence of this specimen together with sequences of all the directly dated and modern specimens. The settings were similar to those used for the leave-one-out analysis. We set a gamma prior (shape = 2; scale = 50,000) on the age of the not directly dated specimens and increased the operator weight of the age estimate to 5. We set a Skygrid tree prior and used a CTMC rate reference prior. Each analysis was run with 50 million generations sampled every 5,000 steps and with the first 5 million generations discarded as burn-in. Convergence and stationarity were inspected in Tracer 1.7 (ESS >200 for all parameters).

In the next step, we reconstructed the phylogeny using all the available sequences (n = 199; ‘joint’ dataset). The analysis was run for 200 million generations and sampled every 20,000 generations. We used a strict clock. This dataset provided strong support for the Skygrid tree prior (Table S9). Guided by the results from the ‘dated’ dataset, uniform prior was used on the clock rate, (2E−8, 2E−6) substitutions/site/year^−1^, with an initial value of 2E−7 substitutions/site/year^–1^. The log-normal prior on the age of each not directly dated specimen was set to correspond to the 95% HPD interval, and the mean was equal to the mean of the age estimate from the individual analysis. This analysis was run in duplicate, and separate runs were checked for convergence and stationarity in Tracer v. 1.7. Trees from both runs were combined using *logcombiner* software from the BEAST 1.10.4 suite with the first 20 million generations discarded as burn-in. The combined trees were summarised, and a Maximum Clade Credibility tree was generated in treeannotator.

Table S9. Model testing for the ‘joint’ dataset using GSS. Details are provided in the ‘Analysis details’ column

| Analysis details | logML (replicates) | Mean logML | 2lnBF |
| --- | --- | --- | --- |
| strict clock, Skygrid tree prior, (2E−8, 2E−6) uniform rate prior | −19871,17 | **−19877,91** | **43,51** |
|  | −19884,64 |  |  |
| strict clock, constants pop. Size tree prior, (2E−8, 2E−6) uniform rate prior | −19899,663 | **−19899,663** |  |
|  | −19899,663 |  |  |

#### 1.8. Demographic analyses

The Bayesian Skyline or Skygrid plot is a powerful tool to reconstruct the past demography of a population but is subjected to assumptions, which can lead to erroneous conclusions if violated, see e.g. Heller et al. (2013). The reconstructed phylogeny suggested that the histories of the common vole populations varied by region, so we did not reconstruct the demographics of the entire dataset. We performed the reconstruction only for all WN lineage specimens from Spain 58; Table S2). These specimens originated from a geographically restricted area and represented a continuous population without recorded lineage turnovers.

The common vole demography was reconstructed using the Bayesian Skyline and SkyGrid methods implemented in BEAST 1.10.4. We fixed the specimen’s ages as the median of the age estimates from the joint phylogeny reconstruction (Table S2) and used the tip dates to calibrate the molecular clock. We used the GTR+I+G, strict clock and either the Bayesian Skyline or Bayesian Skygrid tree priors. We set the Skygrid cut-off to 60 ka as suggested by the joint phylogeny and the number of grid points was set to 50. We ran the BEAST analysis in duplicate for 20 million generations sampled every 2,000 generations. The results were checked for convergence and the log and tree files from two replicates were combined using logcombiner with the first 1,000 trees discarded as burn-in. Bayesian Skygrid and Bayesian Skyline demographic reconstructions were performed in Tracer 1.7.

Both plots (Figures S2 and S3) suggested an increase in the effective female population size (Ne_f_) about 35 ka ago followed by a period of stability and followed by a ca. five-fold decrease in Ne_f_ with a minimum around the Early Holocene (11.7–9 ka ago) followed by a slight increase starting in the Middle Holocene.

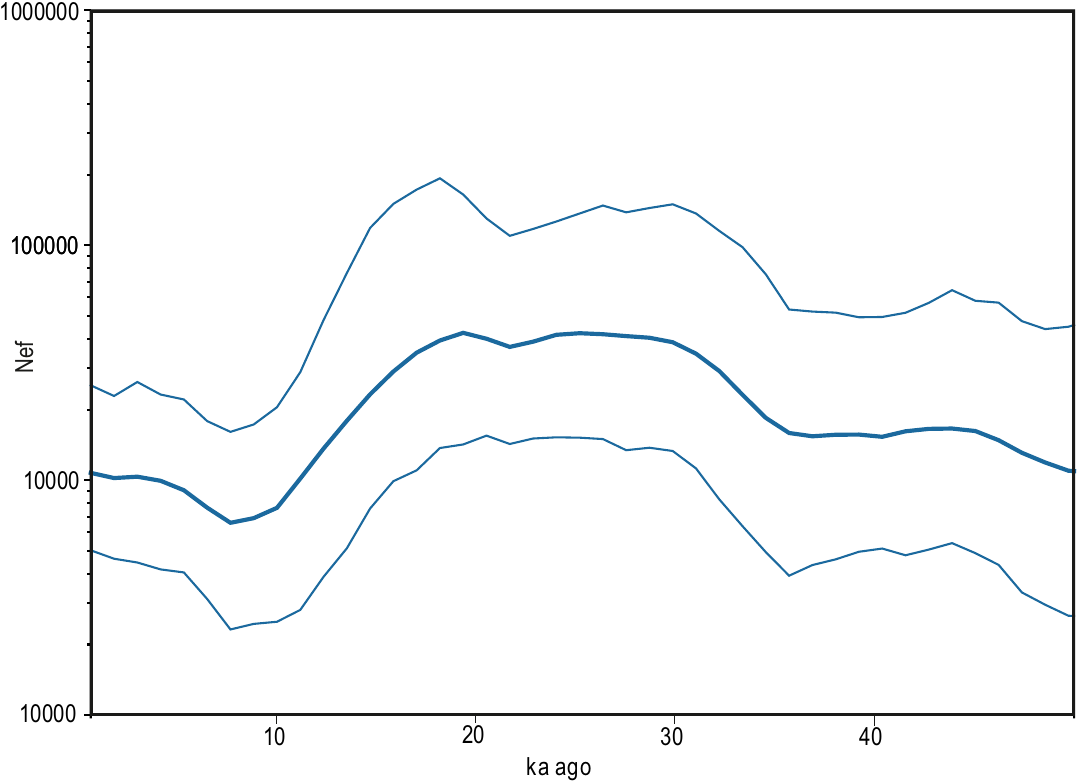

Figure S2. Changes in the effective female population size of common voles from Northern Spain were reconstructed using the Bayesian Skygrid approach implemented in BEAST 1.101.4. The thick solid line denotes the median, and the thin lines mark the limits of the 95% HPD interval.

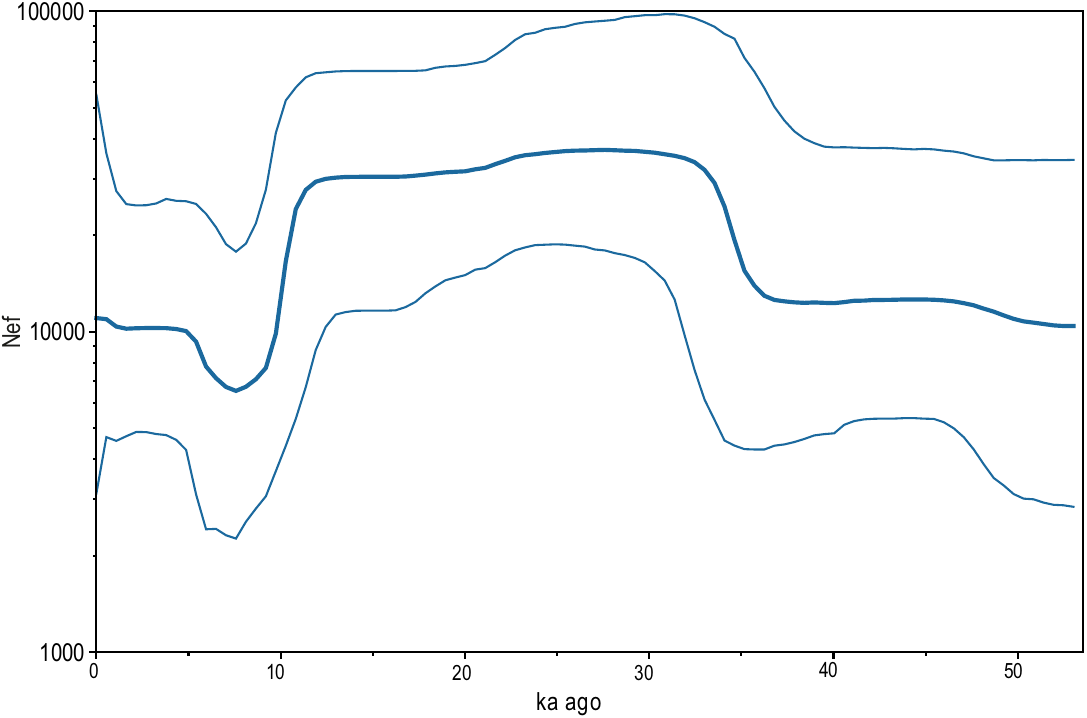

Figure S3. Changes in the effective female population size of common voles from Northern Spain were reconstructed using the Bayesian Skyline approach implemented in BEAST 1.101.4. The thick solid line denotes the median, and the thin lines mark the limits of the 95% HPD interval.

#### 1.9. Radiocarbon dating of vole bones

Helen Fewlass

Vole bones were pre-treated for radiocarbon dating in the Department of Human Evolution at the Max Planck Institute for Evolutionary Anthropology (MPI-EVA, Leipzig, Germany) (lab code: EVA) following the protocol for <100 mg bone samples described in (Fewlass et al., 2019). In brief, the samples were demineralised in 0.5 M HCl at 4°C. Demineralisation was stopped when CO_2_ effervescence had stopped, and the samples were soft (3–23 h). The samples were treated with 0.1 M NaOH for 15 min at room temperature to remove humic acids and then re-acidified in 0.5 M HCl. The samples were rinsed to a neutral pH with ultrapure Milli-Q water between each step. The samples were gelatinised in HCl at pH 3 on a heater block at 70°C until fully solubilised (3–7 h). The resulting gelatine was filtered to remove particles >80 µm (Ezee filters, Elkay labs, UK, cleaned by sonication in Milli-Q water for 20 min) and then ultrafiltered to concentrate the >30 kDa fraction (Sartorius VivaSpin Turbo 15 with 30 kDa molecular weight cut off) with pre-cleaned ultrafilters (Brock et al., 2007). Samples were freeze-dried for 48 h.

The quality of the collagen extracts was assessed based on collagen yield as a percentage of the original bone weight (minimum requirement 1%). The elemental and isotopic ratios of the extracts (~0.5 mg) were measured at the MPI-EVA on a Thermo Finnigan Flash elemental analyser coupled to a Thermo Delta plus XP isotope ratio mass spectrometer. Stable carbon isotope ratios were expressed relative to Vienna PeeDee Belemnite, and stable nitrogen isotope ratios were measured relative to air (atmospheric N2) by using the delta notation (δ) in parts per thousand (‰). Analysis of internal (methionine δ13C = −28.13 ± 0.13 ‰ and δ15N = −6.35 ± 0.03 ‰ (1SD); in-house collagen MRG δ13C = −19.77 ± 0.22 ‰ and δ15N = 5.00 ± 0.06 ‰) and international standards (IAEA-CH-6 sucrose, δ13C = −10.4 ± 0.25 ‰; IAEA-CH-7 polyethylene, δ13C = −32.15 ± 0.14 ‰; IAEA-N-1 ammonium sulphate, δ15N = 0.43 ± 0.04 ‰; IAEA-N-2 ammonium sulphate, δ15N = 20.41 ± 0.2 ‰) indicates an analytical error of 0.2‰ (1σ) for δ13C and δ15N. Collagen extracts were considered suitable for dating where the collagen yield was >1% and elemental values (C: 30%–45%, N: 11%–16%, C:N: 2.9–3.6) fell within established ranges of well-preserved collagen (DeNiro 1985; van Klinken 1999).

When sufficient collagen was extracted (>2 mg), collagen was weighed into tin cups and graphitised using the automated graphitisation equipment (Wacker et al., 2010a) in the Lab of Ion Beam Physics at ETH-Zurich (Switzerland) and dated on a Mini Carbon Dating System (MICADAS) accelerator mass spectrometer (AMS) (Wacker et al., 2010b) (AMS lab code: ETH). When extracted collagen was <2 mg, collagen was weighed into tin cups, combusted to CO2 and measured directly using the gas interface system coupled to the gas ion source of MICADAS (Wacker et al., 2013) as previously described (Fewlass et al., 2018, 2019). Background bones (>50,000 years) of equal size were prepared and dated alongside the vole samples to monitor lab-based contamination. Oxalic acid standards and background collagen samples measured in the same session were used to calculate the age of the samples with BATS software (Wacker et al., 2010a). External errors of 1‰ and 4‰ were propagated in the error calculation of the graphite and gas samples, respectively. Radiocarbon dates were calibrated in OxCal v4.4 (Bronk Ramsey, 2009) by using the IntCal20 (Reimer et al., 2020) calibration curve.

#### 1.10. Additional radiocarbon dates

Mateusz Baca, Adam Nadachowski

Seven radiocarbon dates were obtained from five sites to improve stratigraphic information available for the sites from which the analysed specimens originated. In all the cases, a mix of small mammal bones from the layer of interest was submitted to Poznan Radiocarbon Laboratory (Poznan, Poland). Radiocarbon dates were calibrated in OxCal v4.4 (Bronk Ramsey, 2009) by using the IntCal20 (Reimer et al., 2020) calibration curve (Table S10).

Table S10. Radiocarbon dates obtained to improve stratigraphic information of sites.

| Sample | Country | Site | Layer | Radiocarbon lab ID | Radiocarbon Date | **95% cal BP** | | median | C% | N% | C:N |
| --- | --- | --- | --- | --- | --- | --- | --- | --- | --- | --- | --- |
|  |  |  |  |  |  | **From** | **To** |  |  |  |  |
| MI1262 | Poland | Obłazowa 2 | n/a | Poz-108782 | **33700 ± 700 BP** | 40397 | 36826 | 38547 | 51,22 | 18,40 | 3,25 |
| MI1263 | Poland | Obłazowa 2 | n/a | Poz-108783 | **37100 ± 800 BP** | 42617 | 40736 | 41767 | 48,34 | 17,32 | 3,26 |
| MI1264 | Italy | Grotta del Sambucco | US 5 | Poz-108784 | **19160 ± 120 BP** | 23708 | 22891 | 23094 | 49,56 | 17,71 | 3,26 |
| MI1377 | Slovakia | Muráň 3 | 5 | Poz-122649 | **13510 ± 70 BP** | 16531 | 16056 | 16291 | 47,90 | 17,10 | 3,27 |
| MI2693 | Slovakia | Bišilu | 6 | Poz-122650 | **13290 ± 60 BP** | 16185 | 15767 | 15970 | 49,70 | 17,80 | 3,26 |
| MI2694 | Slovakia | Muráň 3 | 6 | Poz-122651 | **20530 ± 140 BP** | 25104 | 24268 | 24717 | 48,50 | 17,60 | 3,21 |
| MI2696 | Slovakia | Šarkanica |  | Poz-122653 | **21020 ± 140 BP** | 25706 | 25045 | 25379 | 40,60 | 14,60 | 3,24 |

### 2. Description of archaeological/palaeontological sites

Descriptions of most of the sites discussed or mentioned in the text can be found in ‘Appendix A. Supporting Information’ to the paper Baca et al. (2020, *Quaternary Science Reviews*, 233: 106239). Here, only descriptions of new sites from which the studied material originated are presented.

#### 2.1. Bulgaria

Vasil V. Popov

**Cave 16**

43°10 37.31"N 24°4’20.33"E

Cave 16 is a large rock niche, half-filled with sediments, located in Praebalkan region, Karlukovo, north-western Bulgaria. At the entrance, the sediments were exposed by erosion, creating a well-stratified profile over 5 m thick. During pilot archaeological excavations in 1984–1993, the profile was cleared, and its stratigraphy was studied. Fifteen layers were identified (Ferrier, 1994, Popov, 1994, 2000). At present and in the recent geological past, the cave is and has been a nesting place for many petrophilous birds, including owls. This is evidenced by numerous owl pellets on the modern surface of the sediments and the high accumulation of bones in the deposits from layer 15 upwards. Based on the ecological appearance of the stratigraphic small mammal assemblages, the layers formed two distinct groups, namely, upper (layers 9–1) and lower (layers 15–10). Among the layers forming the upper series, particular attention was paid to layer 7, which consists of volcanic ash. It resulted from one of the most powerful volcanic eruptions in southern Italy, known as the Campanian Ignimbrite, dating from ca. 40,000 years BP (Giaccio et al., 2008). The assemblages from the lower part of the profile are characterised by a high proportion of forest and mesophilic species and thermophilic sub-Mediterranean elements; the inhabitants of the steppes have a low share. Quantitative paleoecological reconstructions (Popov, 2014, 2018) suggest that during the formation of the lower part of the sediment sequence (layers 14–10), the temperatures were similar to the modern ones. However, the humidity for most of this period was higher than the present humidity. The climate and vegetation were comparable to those in the mountains of Central Europe. The small mammal assemblages at the top of the profile show a high percentage of species characteristic of open landscapes. The layers of the upper part of the profile (layers 9-1) formed in a cooler climate, especially in the layers deposited before and after the volcanic ash. Winter temperatures were approximately 10 degrees lower than today. The seasonal temperature contrast was significant. The rainfall was less than today. In the age’s context of volcanic ash and based on the ecological appearance of small mammals and quantitative climatic reconstructions, layers 14–10, showing a milder climate, are most likely to be referred to the end of the last interglacial period; layers 10–8 reflect the progressive cooling during the early Last Glacial period (Marine Isotope Stage (MIS) 4–3), corresponding to the time interval between 90,000 and 50,000 years BP; layers 6–1 correlate with the Last Glacial Maximum (LGM).

#### 2.2. Croatia

Jadranka Mauch Lenardić

**Mujina pećina**

43° 33'N 16° 14'E

Mujina pećina is a small, southeast-facing cave located ca. 4.5 km north of Trogir and ca. 16.5 km northwest of Split in middle Dalmatia. It is approximately 10 m deep and 8 m wide. First finds were collected in 1977, and the first test trench was carried out in 1978. Later, it was annually excavated from 1995 until 2003 (Karavanić et al., 2008, 2021). So far, it is the only systematically excavated Middle Palaeolithic site in the Eastern Adriatic with relatively thick homogenous Mousterian deposits in clear stratigraphic context, which has been well-dated chronometrically (Boschian et al*.*, 2017; Rink et al., 2002). Excavation levels followed the natural stratigraphy, and all sediments were sieved. Miracle (2005) presented evidence that humans were the primary accumulation factor for the numerous remains of chamois, ibex, deer, aurochs and bison, whereas the bones of equids and hares were most likely the result of carnivore (e.g. hyena) activity. Cave was not used by humans during summers or winters, when it was probably used as a bear den. In the micromammal sample, representatives of genus *Microtus* prevailed; other small animal taxa suggest open and rocky environment without thick vegetational cover during the deposit sedimentation. Majority of bird taxa also prefer open and rocky habitats, except for *Lagopus lagopus* (boreal) and *Fringilla montifringilla* (tundra) (Mauch Lenardić et al., 2018). Samples analysed in this paper originated from layer B, for which Boschian et al. (2017) quoted that charcoal and pollen of pine, spruce, birch, grassland vegetation and heliophilic shrub joint-pine were found.

Radiocarbon dates from several layers suggest a period between approximately 49 and 39 ka cal BP (Karavanić et al., 2021). For one bone sample from layer B (GrA-9633), the age of 39.2 ka was given (Rink et al., 2002). Meanwhile, Boschian et al. (2017) reported that the radiocarbon age of layer B was 45,250–41,850 years BP.

**Romualdova pećina**

45°7'N 13°42'E

Romualdova pećina (Romuald’s cave) is situated on the southern side of the end part of the Lim Channel near Rovinj in western Istria at the height of 106 m a.s.l. (Malez, 1962). The cave is 110 m long and consists of a long channel which widens on some places and creates elongated chambers with speleothems (Janković et al., 2016; Malez, 1962, 1968; Ruiz-Redondo et al., 2019). Malez noticed ‘bear's polishings’ on the walls during his first visit in 1960, when he also made two test trenches. Later in 1960s and 1970s, systematic excavations were conducted (Malez, 1968, 1978), and rich faunal assemblage was discovered, as well as Upper Palaeolithic lithic tools. In the faunal material, cave bear finds prevailed (Malez, 1987)*.* Komšo (2008) conducted new excavations to revise Malez’s stratigraphy. Aside from Late Pleistocene and Holocene deposits, they also discovered Middle Palaeolithic artefacts and faunal remains. Komšo (2008) and Janković et al. (2016) continued with investigations of Romualdova pećina. They recorded that the layer with several lithic tools below the upper sequences belongs to the Middle Palaeolithic. Furthermore, Ruiz-Redondo et al. (2019) reported that below the topmost Bronze Age layer (3.4–3.2 ka cal BP) was an approximately 1.5 m thick sequence of culturally sterile, natural layers. An underlying 0.2 m thick sequence spanning 34–31.5 ka cal BP yielded Upper Palaeolithic artefacts. Below this was another 0.4 m thick sequence of sterile layers; these layers sealed the Middle Palaeolithic layers, which date to at least 44.3 ka cal BP. A number of red paintings inside the cave were discovered by Komšo in 2010, when he assumed that they are of Palaeolithic age (Ruiz-Redondo et al., 2019). Janković et al. (2016) typologically attributed the lithic material in layers 11 to 13 to the Middle Palaeolithic period for which the age of over 48 ka was obtained. Mauch Lenardić (2013, 2014) analysed small mammal assemblage. She discovered the first remains of lemming in Croatia (*Dicrostonyx* sp.; Mauch Lenardić, 2013) among other arvicolines, which prevailed in small mammal assemblage aside from Chiroptera and Eulipotyphla. The late Pleistocene faunal assemblage of Romualdova Pećina consisted of animals characteristic for more forest environments (e.g. *Clethrionomys* (=*Myodes*) *glareolus*, *Martes martes*, *Felis silvestris*, *Lynx* sp*.*, *Capreolus capreolus*, *Sus scrofa*, *Tetrao tetrix*, *Falco subbuteo*, *Oriolus oriolus*), intermixed with open and rocky areas (*Arvicola amphibius*, *Microtus* ex. gr. *arvalis*/*agrestis*, *Microtus* ex. gr. *subterraneus*/*multiplex*, *Microtus oeconomus*, *Chionomys nivalis*, *Rupicapra* sp., *Equus ferus*, *Perdix perdix*, *Coturnix coturnix*, *Accipiter gentilis*, *Pyrrhocorax* sp*.*). While most of the taxa live in temperate climate and inhabit the vicinity of the cave today, some others do not live in Istria anymore, such as *Dicrostonyx* sp., *Lepus timidus*, *Castor fiber*, *Canis lupus*, *Alces alces*, *Rupicapra* sp., *Lagopus* *lagopus*, and *Pyrrhocorax* sp*.*, indicating the southward movement of the Alpine species during colder periods (Malez, 1968; Mauch Lenardić et al., 2018).

#### 2.3. Czech Republic

Ivan Horáček and Tereza Hadravová

**Bišilu cave**

49^0^ 57' N 14^0^ 6' E, 250 m a.s.l.

The site is located near Tetín, Beroun distr., central Bohemia. A cave entrance in a rocky cliff completely infilled by sedimentary sequence of soil colluvia, limestone debris and loess loam at base. Excavation in 2015-2017 revealed 16 different horizons (with total thickness > 3 m). Deeper part of the section (layers 4b- 8 covering period from 11 ka cal. BP (layer 4b) to > 24 ka cal. BP (data for layer 7a)) was particularly rich in fossils. 28 species of small ground mammals were recorded with total MNI=602. *M. arvalis* appeared in all layers together with *L. anglicus (=gregalis), C. nivalis, A. oeconomus, Lemmus lemmus, Cricetulus migratorius* or *Allactaga major* with peak abundance in layer 6 (16 ka cal BP. - see above).

**Skalice**

490 54'N 140 6'E, 390 m a.s.l.

The cave is located near Měňany, Na Skalici cave, Beroun distr., central Bohemia. This site is a stratigraphical section (>3 m of total thickness) in a sedimentary talus completely infilling a cave entrance in a rocky cliff close to local water spring. Horáček et al. (2002) surveyed the results of the first stage of excavation in 1997. The most recent large-scale excavation was conducted in 2016–2017; analyses are in progress. Layer 7 (from which the items surveyed in this paper originate) represents a base loess with a glacial community dominated by *L. anglicus (gregalis)*, *M. arvalis/agrestis*, *A. oeconomus, C. nivalis*, *Dicrostonyx* sp. and *Ochotona pusilla*.

#### 2.4. France

Aurélien Royer and Loïc Lebreton

**Jovelle**

45°21’37’’N 0°25’48’’E

The cave of Jovelle is located at La Tour Blanche in Dordogne (160 m a.s.l.), Western France, and is constituted as a complex of several loci from distinct Palaeolithic to modern periods. The small mammal materials originated from Diaclase n°2, which is a hyena’s den from the MIS 3. A radiocarbon date (Beta-448744) performed on a bovid metapod showed tooth marks from carnivores, suggesting an occupation older than 43.5 ka. An excavation will be realised soon to investigate the stratigraphic sequence. The main rodent identified in association with *M. arvalis* is *L. anglicus (gregalis)*. Few remains of *A. amphibus*, *D. torquatus,* *A. oeconomus* and *Spermophilus* sp. confirm the glacial period context. They are also associated with more forested taxa, particularly *C. glareolus* and *Apodemus* sp.

**Roc-en-Pail**

47°2’N 0°46’W

Roc-en-Pail is an open-air site located near Chalonnes-sur-Loire, western France, preserved in slope deposits resting on a remnant of a lower fluvial terrace from the Layon River, a small left bank tributary of the Loire River. It was incidentally discovered during winter 1870/1871 (Biaille, 1904) on the margin of a quarry, which was exploiting a limestone hill at the foot of which the Palaeolithic occupations are preserved (Soriano et al., 2021). The excavations of Dr. Gruet during 1940–1950 and 1969 revealed a stratigraphic sequence up to 5 m thick, with seven archaeological layers from the Middle Palaeolithic and one from the Upper Palaeolithic (Gruet, 1945, 1984). New excavation directed by S. Soriano in 2014 and from 2016 to 2018 exposed the top part of the eastern section of the 1969 trench to describe the stratigraphy with further details and establish the chronology. Newly excavated stratigraphic units yielded remains from the Middle Palaeolithic with the exception of the two latest units at the top of the stratigraphic sequence which are still culturally undetermined. Awaiting OSL dating results, the preliminary chronology relies only on geomorphology and characters of faunas and industries. Fluvial deposits at the base of the stratigraphic sequence are attributed to MIS 5, whereas slope deposits accumulated from late MIS 5 to MIS 3. Lithic industries are associated with faunas and microfaunas in all layers; these last were recovered through systematic sieving. DNA analysis highlighted the presence of *Bison schoetensacki* and *Rangifer tarandus* in a MIS 5 layer (Utge et al., 2020). One of the densest occupations can be attributed to the Quina type Mousterian, whereas the latest Middle Palaeolithic occupations present notched and denticulate lithic tools and discoidal debitage.

Among the small vertebrates, small mammals are by far the best represented (more than 90% of the number of remains), followed by birds (3%) and lagomorphs (between (between 1% and 2%). Of the 12 taxa identified, three species are highly abundant (90% of the total NMI), namely, *D. torquatus* (collared lemming), *L. anglicus* (*gregalis)* (narrow-headed vole) and *M. arvalis/agrestis* (field/common vole). Micromammal communities from most of layers exposed in recent excavation reflect a cold and dry climate, contemporary of MIS 4 (Lebreton and Soriano, 2018). The Roc-en-Pail microfauna analysed in this paper was recovered in a late Middle Palaeolithic layer (SU 304) which is considered to be of this age.

#### 2.5. Germany

Sara Rhodes and Nicholas J. Conard

**Geißenklösterle**

48°23’54’’N 9°46’20’’E

Geißenklösterle cave is located in the former Danube Valley approx. 60 m above the valley floor where the Ach River flows today. One of a number of archaeologically important cave sites in the Swabian Jura of southwestern Germany, Geißenklösterle was initially excavated by Eberhard Wagner in 1973. Fieldwork continued under Joachim Hahn from 1974 to 1991 and by Conard et al. from 2000 to 2002 (Conard and Malina 2003), revealing a sequence of deposits spanning the Middle Palaeolithic through to the Holocene. The stratigraphic record at Geißenklösterle is divided into geological horizons (GH) and archaeological horizons (AH), which often but not always correspond to one another. The Middle Palaeolithic deposits correspond to GH 18–23 and AH IV–VIII. Electron spin resonance (ESR) on fossil tooth enamel dates these layers to MIS 5 to the second half of MIS 3 (94 +/- 10 ka and 55 +/- 6 ka) (Richard et al., 2019). Directly overlying the most recent Middle Palaeolithic horizon is a nearly culturally sterile geogenic deposit, GH 17. Radiocarbon and ESR dates place this horizon at ~46 ka and suggest Neanderthals abandoned the site, possibly in response to Heinrich Stadial 5. The Aurignacian occupation documents the arrival of *H. sapiens* in the region as early as 42,500 cal BP (Higham et al., 2012). These groups practiced complex symbolic expression through a combination of portable artworks (i.e. figurines), beads and musical instruments (Conard 2009; Wolf 2013). In total, the Aurignacian period spanned GH 11 through 16 and AH IIn–IIIb.

#### 2.6. Italy

Elisa Luzi

**Grotta del Sambuco**

43°03’N 10°53’E

Grotta del Sambuco is located in central-western Italy (Pianizzoli, Massa Marittima, Tuscany) at 280 m ca. a.s.l. In 2013, the excavation campaigns of the University of Siena revealed a sequence with three late Pleistocene Stratigraphic Units (US). US 4 yielded a rich lithic tools complex (962 pieces) related to final Epigravettian culture (Calattini et al., 2016) and dated at 13.615±75 BP (uncalibrated). US 5 was extremely rich in small mammals, with scarce lithic industries and numerous charcoals. In 2018, a human femur was recovered from this unit. US 6 returned abundant small mammals, scarce lithic tools and large mammal rests (horse, wolf and red deer), and marine malacofauna (*Bittium*) (Calattini et al., 2019). US 6 was dated at 23.632 ± 150 BP (uncalibrated). The small mammal assemblages of US 5 and US 6 were dominated by the common vole *M.* *arvalis* (> 90% in both Units) (Luzi et al., 2020).

#### 2.7. Poland

Anna Lemanik and Adam Nadachowski

**Deszczowa cave**

50°34'N 19°31'E

Deszczowa cave, located in Kraków – Częstochowa Upland, southern Poland, was excavated in 1989–1997. Four sedimentary series and 11 layers were distinguished within the exposed profile. The sedimentation occurred from the late Saalian, probably to the beginning of the Eemian Interglacial (MIS 6 to MIS 5e), and after an erosional hiatus during MIS 3 and MIS 2 and the Holocene (Cyrek et al., 2000; Nadachowski et al., 2009; Krajcarz and Madeyska, 2010). The deepest layers (I, II and III) filled the narrow fissure of the cave bottom and were archaeologically sterile (Cyrek et al., 2000). The small mammal fauna from layers I, II and III includes the characteristic taxon *M. oeconomus/malei* (Nadachowski et al., 2009), also present in Biśnik Cave, which is indicative in Poland for the late Middle Pleistocene (MIS 6). Layers I and II lay at the rocky bottom, and layer III fills up spaces between large limestone blocs situated below generation of flowstone dated by Th/U method to ca. 125 ka (+18 ka; - 20 ka). The rocky bottom of the cave (flowstone) was dated at ca. 210–217 ka (error ca. +ca. 40 ka / – ca. 34 ka) and some parts at > 350 ka. Therefore, layers I, II and III are older than Eemian but probably younger than MIS 8 (Saalian Complex). Dates (AMS, 14C) are available for the upperpart of the profile (Nadachowski et al., 2009; Lorenc, 2013). For a very short stratigraphic description, see Klimowicz et al. (2016).

**Obłazowa 2**

49°25'48"N 20°09'36"E

Obłazowa 2 site is located in Obłazowa Rock, 50 m from the main entrance to the Obłazowa Cave, Nowa Biała, Western Carpathians, southern Poland. It is an artificial shelter, approximately 7 m long and 3 m wide, with a large opening facing east. In the 1920s and 1930s, the eastern part of Obłazowa Rock was exploited as a quarry and an artificial cave was created. The shelter was established in a natural fissure (with an original width of approx. 0.5 m) developed at the contact zone between the crinoid limestone and red nodular limestone and filled with a large amount of sharp-edged limestone rubble and loamy clay which is intense brownish-red in colour. The fossil-bearing sediment contained regularly distributed sub-fossil bones and teeth. The rich fauna of vertebrates consisted of rare fishes, amphibians and reptiles and abundant remains of birds (38 taxa) and mammals (33 species) (total NISP=9490; MNI=4636) (Nadachowski et al., 1993). The faunal composition indicates a park-steppe and/or forest-steppe biomes, abundant in humid environments. The fauna was dated to the middle part of MIS 3. One date was obtained from a fox mandible: OxA-3696, 33,430 ± 1230 BP) and two further dates are presented in Table S10.

#### 2.8. Romania

Alexandru Petculescu

**Hoților cave**

44°53'47.17"N 22°25'43.32"E, approx. 200 m a.s.l.

Hoților cave is located on the right side of the Cerna river, near Baile Herculane locality, Caraș-Severin County. This cave has been popular, especially from the biospeological point of view, since 1898. In 1954, the Institute of Archaeology from Bucharest carried out extensive archaeological investigations in this cave and found Palaeolithic and Neolithic traces as well lithic industry. It is a small cave but with a wide opening and water sources in proximity, favourable for human habitation.

E. Terzea (1971, 1986) analysed the association of the following small mammals in this cave: *Talpa europaea, Crocidura leucodon, Sorex minutus, Sorex araneus, Citellus* sp*., Dryomys nitedula, Muscardinus avellanarius, Sicista* cf*. subtilis, Spalax* sp*., Apodemus sylvaticus, Cricetulus migratorius, Cricetus cricetus, C. glareolus, A. amphibius, C. nivalis, A. oeconomus, L. gregalis, M. arvalis, M. agrestris* and *M. subterraneus*.

The *M. arvalis* sample originated from layer 4 (Aurignacian) (Terzea, 1986), which has a mixed small mammal fauna association showing a mild temperate climate period favourable for forest and steppe areas.

**Muierilor cave**

45°11'31.78"N 23°45'14.07"E, 650 m a.s.l.

Muierilor cave is an extensive cave system and a popular show cave in Romania. It is located in the southern part of the Parâng mountains, near Baia de Fier city, Gorj County, on the right side of the Galbenului Valley. This cave is a well-known archaeological, anthropological and palaeontological site which holds impressive Marine Isotope Stage 3 fossil association (Doboș et al., 2010).

The fossil bone accumulation was identified in the 1870s; the first systematic excavations in Muierilor cave started in 1929 and continued during the 1950s. The detailed large mammal association from this cave was first determined by Bombiță (1954) and included *Ursus spelaeus, P. spelaea, Panthera pardus, Lynx lynx, F. silvestris, C. lupus, Crocuta crocuta spelaea, Vulpes vulpes, Gulo gulo, Lutra lutra, M. martes, Saiga tatarica, Capra ibex, Rupicapra rupicapra, Cervus elaphus, Megaloceros giganteus, Bison priscus, Bos primigenius, A. alces, Equus caballus, Rhinoceros tichorhinus* and *Mammuthus primigenius.*

Holocene human remains were found close to the entrance area, whereas the Upper Palaeolithic human remains were found much deeper in the cave, in the Mousterian Passage (Nicolăescu-Plopșor et al., 1957; Păunescu et al., 1982) which are one of the most important discoveries in Muierilor cave until now, the remains of one of the earliest anatomically modern humans in Europe, directly dated to ~35 ka cal BP (Soficaru et al. 2006).

Part of the Muierilor cave, Bears Passage hosts a large amount of fossil remains, which determined us to establish here, starting from 2014, the new palaeontological excavation composed from nine squares, 1 sqm each, who are transposed transversally on the Bears Passage. The trench was designed to intercept the entire width of the Bears Passage on the excavation site to reveal the palaeohydrological conditions of the sediment and fossils accumulation.

The maximum depth resulting from our excavation reaches 260 cm in the central part with ten sedimentary layers. The first 30 cm is rich in fossil remains dominated by *U. spelaeus* (69.5%), followed by *C. lupus, Crocuta crocuta spelaea, V. vulpes*, *P. spelaea* and small mammals (Mirea et al., 2020).

The small mammal association from the new excavation includes the following eight species: *M. arvalis* (40%), *Lagurus lagurus, C. nivalis, Eolagurus luteus, A. amphibius, C. glareolus, C. cricetus* and *S. araneus*, but the work is still in progress.

The *M. arvalis* sample used for this paper has been provided by the first layer from the surface (Fig. 1, L1), dated between 14.7 ka (the base of a stalagmite grown on the cave floor, U/th) and 39.6 ka BP (C14 on fossil bone) calibrated age (Mirea et al., 2020).

**Stoieni cave**

44°58'51.6"N 22°36'14.9"E, 700 m a.s.l.

This palaeontological site is relatively recently discovered. First excavations were carried out since 2009 by Emil Racovita Speleological Institute team, currently reaching a depth of 2 m and 2 sqm. This cave is situated close to Isverna village, Mehedinți Plateau, a vast karstic region in southwestern Romania. From the first four levels (0–60 cm), 14 species of small mammals were identified from over 500 individuals. The list of large mammals in this cave includes *C. ibex, C. lupus, Ursus arctos, U. spelaeus, V. vulpes, Meles meles*, *Lepus* sp., *Felis* sp., *Ovis* sp. and Aves.

The lowermost sampled layer (level 4, 45–60 cm) from where the material for this study was selected, is characterised by a relatively high abundance of *C. nivalis* and *M. arvalis*, indicating a rough climate and the presence of dwarf bush associations. The extremely low abundance of dormouse (*Glis glis*) reveals a lack of forest cover and supports the scenario of a cold and dry environment during this stage. The second and third depth intervals (15–30 and 30–45 cm) are dominated by a large population of dormouse, which is a typical forest inhabitant, whereas the numbers of *C. nivalis* and *M. arvalis* are decreasing, indicating an extensive forest cover in a dominant temperate to cold climate (Petculescu, 2013).

The uppermost depth interval (0–15 cm) is again defined by a high abundance of *C. nivalis*, indicating the presence of a cooling event. In addition, the presence of a small number of *C. glareolus* and *Spalax leucodon* counts for a steppic environment, whereas *A. flavicollis* and the numerous *G. glis* population, both tree-climbing taxa, indicate that tall vegetation was still abundant during this stage. Conversely, the large number of dormice can be explained by the fact that this species hibernates in cracks and caves, and the prolongation of the cold season can easily lead to a high mortality rate because of decreased feeding possibilities (Kryštufek, 2008).

#### 2.9. Serbia

Zoran Marković

**Gradašnica cave**

N 44° 27' 45" E 22º 13' 42"

Gradašnica cave is located on the west side of Veliki Greben and Veliki Strnjak, nearby the Miroč village, above the Danube gorge. The cave’s monumental entrance is 35 m high and 15 m wide. Its altitude is approximately 380 m above the sea level. The main channel caved in Jurassic limestone is approximately 300 m long. The cave belongs to the spring-type of caves (Petrović, 1976). First palaeontological investigation of this cave started in 1998 (Marković, 2008). High water level throughout the year and the reduced accessibility to all parts (too low ceilings and too narrow passages) conditioned a limited choice of sediment collection sites. Approximately 50 kg of sediments were collected from the left ‘dead-end’ passage on the entrance of the cave. This passage, which is very narrow, contains a wider area of approximately 2.5 m^2^ where the sediments were neither disturbed by water nor by human activity. The porous clay sediment, which covers the limestone slope, was nowhere thicker than 15 cm. Variations in its lithological composition were undetectable, and layers were not recognised. After sieving the sediment, the remains of animals from different classes are sorted: Amphibia, Reptilia*,* Aves and Mammalia. The mammal remains were the most numerous, representing approximately 80% out of 2500 items: *Rhinolophus hipposideros, Rhinolophus ferrumequinum, Myotis myotis, Таlра europea, S. araneus, Neomys fodiens, C. leucodon, Spermophilus citelus, Mesocricetus newtoni, C. migratorius, C. cricetus, C. glareolus, A.* *amphibius, M. arvalis, M. subterraneus, C. nivаlis, Nannospalax leucodon, G. glis, Muscardinus avelanarius, S. subtilis, A. flavicollis/А. sylvaticus, Rattus* *rattus, Ochotona pusilla, Lepus* sp., *V. vulpes*, *M. martes*, *Ursus* sp., Bovidae sp.. The remains were accumulated in a place which is certainly not the habitat of any of these species. The bones did not bear any signs of being transported by water on any other geomorphological agents. Therefore, the most probable means of transporting and accumulation of this material is by the means of predators. Although no recent material was discovered, such a cave is a natural habitat for some of the owl species in the past and occasionally for some members of carnivora (*Mustela nivalis*, *V. vulpes* and *Martes* sp.). Analysis of the pattern of skull and mandible breakage showed that most remains were part of owl prey (Andrews, 1990). Some of the species are extinct in the area at present (*O. pusilla*, *M. newtoni*, *C. migratorius*, *S. subtilis*), whereas the others are still present (Petrov, 1992). The fossil *Rodentia* and *Lagomorpha* fauna from the Gradašnica cave sediments may be characterised by a mixture of steppe and forest faunal elements, with several species belonging to other biotopes, such as forest-steppe, mountain meadows, stony mountain peaks and wet forest clearings (Marković, 2008). None of the species found in the fossil material is extinct.

**Vrelska pećina**

N 43° 12' 53" E 22° 18' 50"

Vrelska pećina cave is located in Bela Palanka town centre (east Serbia). The cave was explored several times since 1986, primarily in search of the water supply purpose. Blasting destroyed much of the cave interior and in many places disturbed the natural sediment relationship (Petrović, 1976). During 1990, several samples (mostly from yellowish gravelly-silty and porous clay sediment 6 m from the cave entrance) resulted from the extremely abundant assemblage of fossil remains: mammals (Mammalia), birds (Aves), reptiles (Reptilia), amphibians (Amphibia), fish (Pisces) and crustaceans (Crustacea) (Marković & Pavlović, 1991). The generally well-preserved remains allowed the identification of the following species: *Sorex* sp., *S. araneus*, *S. minutus*, *Neomys* sp., *C. leucodon*, *Rhinolophus* sp., *R. hipposideros*, *R. mehelyi*, *M. myotis*, *Lepus europeus*, *O. pusilla*, *Spermophilus citellus*, *S. subtilis*, *N. leucadon*, *G. glis*, *A. sylvaticus*, *Mesocricetus newtoni*, *C. migratorius*, *C. cricetus*, *C. glareolus*, *L. lagurus*, *A. terrestris*, *C. nivalis*, *M. arvalis*, *M. agrestis* and *M. subterraneus*. Remains of large mammals were sporadic in the central cave area: *Equus* sp., *B. priscus, B. primigenius*, *C. capreolus, Ursus* sp*.,* *F. silvestris, Lynx* sp. and *Canis* sp. The composition of the mammal fauna suggests that the remains were deposited in the late Upper Pleistocene.

#### 2.10. Slovakia

Ivan Horáček and Tereza Hadravová

**Dzerava skala cave**

480 30'N 170 19'E, 410 m a.s.l.

The site is located in Malé Karpaty Mts., southwest Slovakia. A spacious cave entrance has been recently excavated in 2002–2003 by Kaminská et al. (2005). With nine layers, the section and abundant faunal remains (small mammals MNI=2013) documented faunal development from ca 50 ka to 25 ka BP (18 ^14^C data from 28,865 to 53,305 cal BP). The samples analysed in this paper originated from layers 3 and 4, representing early MIS2 to late MIS3 (two ^14^C data from 28,265 to 31,033 cal BP). The mammal fauna (MNI 480 and 369) was dominated by *M. arvalis* (27.1%), followed by *L. anglicus (=gregalis), Dicrostonyx* sp., *A. oeconomus, C. nivalis, O. pusilla, Lepus* sp., *S. araneus* and 16 species with recedent representation. For details, see Horáček (2005).

**Nový Mt. - Horní cave III,**

49°11' 20°14'E, 1791 m a.s.l.

The site is located in High Tatras Mts., North Slovakia**.**The items surveyed in this paper come from the middle layers (nos. 3 and 4) of a cave infill which consists of the cryogenic rubble almost without any loamy material – maximum thickness of the bottom sediment was less than 1 m (Sekyra, 1954). Samples collected in 1975 by V. Ložek, the other abundant material reported by Schaefer (1975) who provided a ^14^C dating 30 ky. Comp. also Horáček et al. (2015).

**Šarkanica cave**

480 43 N 190 58'E, 925 m a.s.l.

The site is located near Tisovec, Muráňská planina Mts., South Slovakia. Enormously rich bone deposit (MNI in total 6586) originated from three nest sites of *Nyctea scandiaca* in the surface layer of loess loams infilling in a spacious cave entrance. The material was collected in 1983 by Jan Obuch, who also provided a detailed site description (Obuch, 2006). Four ^14^C data varied from 17970 BP to 21400 BP (Kaizer et al., 2018).A smaller part of the material (MNI=795) deposited in Zoology coll. Charles Univ. Prague (from which the samples surveyed in this paper were obtained) is composed of the following (MNI): 4 *S. araneus*, 16 *Sorex tundrensis*, one *Neomys anomalus*, one *Dicrostonyx* sp., seven *C. glareolus*, six *A. amphibius*, 58 *C. nivalis*, 598 *L. anglicus (gregalis)*, 47 *A. oeconomus*, 32 *M. agrestis*, 22 *M. arvalis*, one *O. pusilla*, one *Mustela* cf. *nivalis.* These samples undoubtedly represent a pure community of the LGM stage.

**Peskö cave**

48°29’N 20°21É, 210 m a.s.l.

The site is located near Bretka, Rimava basin, south Slovakia. More than 4 m deep stratigraphic section at the entrance of a small cave situated in a limestone cliff over a bank of Muráň river about 1 km N of village Bretka at the northern margin of Rimava basin, the northernmost outskirts of Pannonian lowland. The excavation was conducted from 1984 to 1987. For detailed information, see Ložek et al. (1989). The section revealed a sequence of 17 horizons carefully sampled for mollusc and vertebrate fossil remains (20–50 kg of sediments per layer). The upper part of the sequence (layers 1–7b, roughly a half of total sediment thickness) contained a series of scree and soil colluvia with rich mollusc and vertebrate communities of a high diversity, including demanding woodland and open ground elements (small mammals MNI 373, 25 spp., bats MNI 41, 17 spp.). It represents the late glacial and early Holocene development. The loess layer at its base (8) reveals discordant bordering with the overlying layer 7b and particularly with the uppermost member of basal sequence (9a). It indicates an eroding event (supposedly associated with the destruction of the cave entrance during LGM) and a considerable time gap in the sedimentary record. The underlying sequence (layers 9–13) of the clayed coarse scree was characterised by dispersed terra rossa conglomerates, poor mollusc remains and rich small mammals (MNI 399, 14 spp., bats MNI 11, 7 spp.). It shows glacial assemblages dominated by *L. anglicus* and *M. arvalis*, which prevailed in the deepest layers (11–13) where the community included *M. agrestis, A. oeconomus, C. nivalis, S. subtilis, C. cricetus*, *Allactaga major* and *L. lagurus*. Radiocarbon dates from this area varied from 36557 ka cal BP (layer 9) to 45417, 45548 and 54027 ka cal BP (layer 12), whereas those from the upper part of the section (layer 4) were 11263 and 14973 ka cal BP.

#### 2.11. Spain

Xabier Murelaga

**Artazu VII**

43°4’26’’N - 2°31’49’’E

Artazu VII site is a fossiliferous deposit, which was accidentally discovered in 2012 after a following basting in Kobate Quarry, Arrasate, Gipuzkoa, 70 km to the southwest of Donostia-San Sebastián, northern Spain and one year later, in 2013, excavated in an emergency excavation by Mari Jose Iriarte and Alvaro Arrizabalaga (Suarez-Bilbao et al, 2016, 2017 and 2018). The site is located at 351 m above sea level. Although a large part of the site was destroyed during the blasting, it was possible to collect some in situ material (Suárez-Bilbao et al., 2016). In the structure of the preserved site, three different areas, but no levels, were differentiated, named, from top to bottom, Upper Ledge, Lower Ledge and Chamber. In total 50 taxa have been identified, 24 of them belonging to small vertebrates (13 mammals, seven amphibians and four reptiles), 14 to large mammals (four ungulates and ten carnivores) and 12 to avifauna. The rodents association includes the following seven taxa: *A. sylvaticus-flavicollis*, *A. amphibiu*s, *A. sapidus*, *M. agrestis*, *M. arvalis*, *M. (Terricola)* sp. and *P. lenki* (Suárez-Bilbao et al. 2017). A humid habitat with a predominance of grassland and broadleaf forests, with some practically permanent water sources, has been inferred on the basis of faunal composition. The absence of species characteristic of cold climates and the large number of indicators of a relatively warm climate (e.g., the presence of the genus *Apodemus*) show that the atmospheric temperature was relatively similar to present conditions (Suarez-Bilbao et al, 2016, 2017 and 2018). The AAR dates of 98.4 ka and 88.5 ka, obtained for bone samples from Layers L and K in the Lower Ledge, suggest an attribution of Artazu VII to the first half of the Late Pleistocene, in the MIS 5c substage (Suarez-Bilbao et al, 2016, 2017 and 2018). Artazu VII is one of the few Pleistocene sites belonging to MIS 5c in the Iberian Peninsula.

#### 2.12. Ukraine

Bogdan Ridush

**Perlyna cave**

48°16'N 23°37'46"E

Perlyna is a small cave, located 1.5 km to the north from Mala Uholka Village of Tiachiv district in Zakarpattia region of western Ukraine. Together with four other caves, it is a part of a single fossil hydrogeological system of the Viv Cliff. Perlyna cave (which means ‘a pearl cave’), is only 36 m in total length, with an amplitude of +6.0 m. It consists of one ascending gallery nearly 1–3 m wide and 1–2 m high with a short and narrow side pass. Its morphology and bedding inside the thick-layered limestone, mainly along the bedding surfaces, indicate their ancient artesian genesis (Ridush, 2012). The cave morphology is of the microclimatic type of a ‘warm bag’. Upward from the entrance is a corridor which starts with a low gallery with an angled transversal profile. Then, we entered a rising room with one wall covered with flowstone. The cavity continues downwards, but it is filled with sediments, including debris, bones, rare inclusions of quartz, jade and sandstone pebbles. Fossil bones occur in debris-loamy sediments (approximately 20–30 cm thick).

First bone remains were collected in 1962. In 1963–1965, the excavations were provided by the Zoological Institute of the Academy of Science of the Ukrainian SSR and by the Zoological Department of the University of Uzhgorod (Bachynsky and Chernysh, 1965; Koliushev, 1966). Some collections were provided in 1989–1993 (Krochko et al., 1993). According to G.O. Bachynskyi, the cave bear (*U. spelaeus*) remains prevailed in the cave, along with bones of martens (*M. martes*), badgers (*Meles meles*), foxes (*V. vulpes*), wild cat (*F. silvestris*), reindeer (*R. tarandus*), bison (*B. priscus*) and other small vertebrates (Bachynsky, 1970). Approximately 1100 bones belonging to 18 cave bear individuals and other species were excavated during the 1960s (Bachynsky and Chernysh, 1965; Tatarinov and Bachynskyi, 1968).

We have surveyed the cave since 2006. In 2017–2018, regular excavations were provided. The top layer, which is bone-bearing, was highly dug and mixed. Some bones were included in the flowstone. The roots of trees were present in the cave floor. Bones were also present in loamy sediments of the upper chamber above the flowstone. Skeletal elements are represented mainly with metapodials, phalanxes and teeth, sometimes with broken long bones. Among megafauna species, the most numerous are bears, including *U. arctos, U.* cf*. deningeri, U.* cf. *suessenbornensis* and *U. ingressus*. Rare species include *Panthera spelaea*, *Lynx lynx*, *F. silvestris* and *C.* *lupus* (Ridush and Popiuk, 2019). The alone radiocarbon date is given for the bone of *U. ingressus* (45,700 +2,500/-1,900, VERA 3736) (Ridush, 2014).

### 3. References

Andrews, P. (1990). Owls, Caves and Fossils - Natural History Museum Publication, London, 231 pp.

Baca, M., Popović. D., Baca, K., Lemanik, A., Doan, K., Horáček, I., López-Garcia, J. M., Bañuls-Cardona, S., Pazonyi, P., Desclaux, E., Crégut-Bonnoure, E., Berto, C., Mauch Lenardić, J., Miękina, B., Murelaga, X., Cuenca-Bescós, G., Krajcarz, M., Marković, Z., Petculescu, A., Wilczyński, J., Knul, M. V., Stewart, J. R., Nadachowski, A. (2020). Diverse responses of common vole (*Microtus arvalis*) populations to Late Glacial and Early Holocene climate changes – Evidence from ancient DNA. *Quaternary Science Reviews*, 233(106239).

Baca, M., Popović, D., Lemanik, A., Baca, K., Horáček, I., & Nadachowski, A. (2019). Highly divergent lineage of narrow-headed vole from the Late Pleistocene Europe. *Scientific Reports*, *9*(1), 17799. doi: 10.1038/s41598-019-53937-1

Bachynsky, G. A. (1970). Tafonomichna kharakterystyka mistseznakhodzhen vykopnych kherebetnykh v karstovykh pecherakh Ukrainy [Taphonomic characteristics of the findings of the fossil vertebrates in the caves of Ukraine]. *Fizychna Geografia ta Geomorfologia,* 153–160.

Bachynsky, G. O., Chernysh, I. V. (1965). Nove pecherne mistseznakhodzhenia vykopnykh khrebetnykh v Ukrainskykh Karpatakh [New cave site of fossil Vertebrates in the Ukranian Carpathians]. *Dopovidi Akademii Nauk Ukraiskoi RSR,* 12, 1631–1633.

Baele, G., Lemey, P., Suchard, M. A. (2016). Genealogical working distributions for Bayesian model testing with phylogenetic uncertainty. *Systematic Biology,* 65, 250–264.

Biaille, M. (1904). Silex et ossements trouvés au confluent de la Loire et du Layon , in Compte-rendu de la 32e session, Angers, 1903, Seconde partie, Notes et mémoires, Paris: Association française pour l’avancement des sciences, 862-863.

Bombiță, G., (1954). “Mammals from Baia de Fier caves. Results from paleontological excavations made in 1951” Scietific report*. Secțiunea de Științe Biologice, Agronomie, Geologice si Geografice,* 6, 253–99.

Boschian, G., Gerometta, K., Ellwood, B. B., Karavanić, I. (2017). Late Neandertals in Dalmatia: Site formation processes, chronology, climate change and human activity at Mujina Pećina, Croatia. *Quaternary International* 450, 12–35. https://dx.doi.org/10.1016/j.quaint.2016.09.066.

Brock, F., Bronk Ramsey, C., Higham, T. (2007). Quality assurance of ultrafiltered bone dating. *Radiocarbon,* 49, 187–92.

Bronk Ramsey, C. (2009). Bayesian analysis of radiocarbon dates. *Radiocarbon,* 51: 337–360.

Calattini, M., Galiberti, A., Tessaro, C. (2016). Massa Marittima (GR). Grotta del Sambuco: indagini 2015, Notiziario della Soprintendenza per i Beni Archeologici della Toscana 11. Firenze.

Calattini, M., Galiberti, A., Tessaro, C. (2019). La Grotta del Sambuco (Massa Marittima, GR): Risultati delle ultime ricerche. *Boll. di Archeol*, Online X, 27–36.

Conard, N. J. (2009). Female figurine from the basal Aurignacian of Hohle Fels Cave in southwestern Germany. *Nature,* 459**,** 248–252.

Conard, N. J., Malina, M. (2003). Abschließende Ausgrabungen im Geißenklösterle bei Blaubeuren, Alb-Donau-Kreis. In: Archäologische Ausgrabungen in Baden- Württemberg 2002. Theiss Verlag, Stuttgart.

Cyrek, K., Nadachowski, A., Madeyska, T., Bocheński, Z., Tomek, T., Wojtal, P., Miękina, B., Lipecki, G., Garapich, A., Rzebik-Kowalska, B., Stworzewicz, E., Wolsan, M., Godawa, J., Kościów, R., Fostowicz-Frelik, L., Szyndlar, Z. (2000). Excavation in the Deszczowa Cave (Kroczyckie Rocks, Częstochowa Upland, Central Poland). *Folia Quaternaria,* 71, 5-84.

DeNiro, M. J. (1985). Postmortem preservation and alteration of in vivo bone collagen isotope ratios in relation to palaeodietary reconstruction. *Nature,* 317(6040), 806-9.

Doboș, A., Soficaru, A., Trinkaus, E., (2010). The Prehistory and Paleontology of the Peștera Muierii, Romania. Liege.

Duchene, S., Lemey, P., Stadler, T., Ho, S. Y. W., Duchene, D. A., Dhanasekaran, V., Baele, G. (2020). Bayesian evaluation of temporal signal in measurably evolving populations. *Molecular Biology and Evolution*, 37, 3363–3379. https://doi.org/10.1093/molbe v/msaa163.

Ferreira, M. A., Suchard, M. A. (2008). Bayesian analysis of elapsed times in continuous-time Markov chains. *Canadian Journal of Statistics,* 36, 355–368.

Ferrier, C. (1994). Le context environmental du peuplement paleolithique de Bulgarie du Nort. Le karst de Karlukovo et ses depots. These. L'Universite Bordeuax I. 451 p.

Fewlass, H., Talamo, S., Tuna, T., Fagault, Y., Kromer, B., Hoffmann, H., Pangrazzi, C., Hublin, J. J., Bard, E. (2018). Size matters: Radiocarbon dates of <200 μg ancient collagen samples with AixMICADAS and its gas ion source, *Radiocarbon,* 60, 425–439.

Fewlass, H., Tuna, T., Fagault, Y., Hublin, J. J., Kromer, B., Bard, E., Talamo, S. (2019). Pretreatment and gaseous radiocarbon dating of 40–100 mg archaeological bone. *Scientific Reports,* 9, 1–11.

Gansauge, M. T., Aximu-Petri, A., Nagel, S., & Meyer, M. (2020). Manual and automated preparation of single-stranded DNA libraries for the sequencing of DNA from ancient biological remains and other sources of highly degraded DNA. *Nature Protocols*, *15*(8), 2279–2300. doi: 10.1038/s41596-020-0338-0

García, J. T., Domínguez-Villaseñor, J., Alda, F., Calero-Riestra, M., Pérez Olea, P., Fargallo, J. A., Martínez-Padilla, J., Herranz, J., Oñate, J.J., Santamaría, A., Motro, Y., Attie, C., Bretagnolle, V., Delibes, J., & Viñuela, J. (2020). A complex scenario of glacial survival in Mediterranean and continental refugia of a temperate continental vole species (*Microtus arvalis*) in Europe. *Journal of Zoological Systematics and Evolutionary Research*, 58, 459–474.

Giaccio, B., Isaia, R., Fedele, F. G., Di Canzio, E., Hoffecker, J. F., Ronchitelli, A., Sinitsyn, A. A., Anikovich, M., Lisitsyn, S. N., Popov, V. V. (2008). The Campanian Ignimbrite and Codola tephra layers: two temporal/stratigraphic markers for the Early Upper Palaeolithic in southern Italy and eastern Europe. *Journal of Volcanology and Geothermal Research,* 177, 208–226.

Gruet, M. (1945). Séance du 25 janvier 1945. Correspondance, *Bulletin de la Société préhistorique de France,* 42(1), 3-4.

Gruet, M. (1984). L’apport de deux sites angevins à la chronologie des terrasses fluviales : Roc-en-Pail en Chalonnes sur Loire et Port- Launay sur la Sarthe. *Bulletin de l’Association française pour l’étude du quaternaire,* 21(1), 13-18.

Heller, R., Chikhi, L., & Siegismund, H.R. (2013). The Confounding Effect of Population Structure on Bayesian Skyline Plot Inferences of Demographic History. *PLoS ONE*, 8(5), e62992.

Higham, T., Basell, L., Jacobi, R., Wood, R., Ramsey, C. B., Conard, N. J. (2012). Testing models for the beginnings of the Aurignacian and the advent of figurative art. and music: the radiocarbon chronology of Geißenklösterle. *Journal of Human Evolution,* 62(6), 664-676. https://doi.org/10.1016/j.jhevol.2012.03.003.

Horáček, I. (2005). Small vertebrates in the Weichselian series in Dzeravá skala cave: list of the samples and a brief summary. Pleistocene Environments and Archaelogy of the Dzerava skala Cave, Lesser Carpathians, Slovakia, 157-167.

Horáček, I., Ložek, V., Svoboda, J., Šajnerová, A. (2002) Environmentand Late Paleolithic/Mesolithic settlement of the karstic areas. Pp. 313-344 In: Svoboda J. (ed): Prehistoric caves – Catalogues, documents, studies. Archeol. Inst AV CR, Brno 407 pp. (in Czech).

Horáček, I., Ložek, V., Knitlová, M., Juřičková, L. (2015). Darkness under candlestick: glacial refugia on mountain glaciers. Forgotten Times and Spaces: New Perspectives in Paleoanthropological, Paleoethnological and Archeological Studies. Brno: *Masaryk University*, 363-377.

Horn, S. (2012) Target enrichment via DNA hybridization capture. Ancient DNA. Methods and Protocols. (ed. by B. Shapiro and M. Hofreiter), pp. 189–95. Humana Press, Totowa, NJ.

Janković, I., Komšo, D., Ahern, J. C. M., Becker, R., Gerometta, K., Mihelić, S., Zubčić, K. (2016). Arhaeološka istraživanja u Limskom kanalu 2014. i 2015. Lokaliteti Romualdova pećina i Abri Kontija 002, Pećina kod Rovinjskog Sela, Lim 001 i podvodni pregled Limskog kanala (Archaeological investigation of the Lim Channel in 2014 and 2015 at Romuald's Cave, Abri Kontija 002, Pećina Cave near Rovinjsko Selo, Lim 001 and an underwater survey of the Lim Channel). *Historia archaeologica,* 46, 5–23. [in Croatian and English].

Kaizer, J., Obuch, J., Kontuľ, I., Šivo, A., Richtáriková, M., Čech, P., Povinec, P. P. (2018) Methods of Radiocarbon Determination in wine and bone samples by Gas Proportional Counting Technique. *Radiocarbon,* 1-11.

Kaminská, L., Kozłowski, J. K., Svoboda, J. (2005). Pleistocene environments and archaeology of the Dzerava skala Cave, Lesser Carpathians, Slovakia. Polish Academy of Arts and Sciences, Kraków.

Karavanić, I., Miracle, P. T., Culiberg, M., Kurtanjek, D., Zupanič, J., Golubić, V., Paunović, M., Mauch Lenardić, J., Malez, V., Šošić, R., Janković, I., Smith, F. H. (2008). The Middle Paleolithic from Mujina pećina, Dalmatia, Croatia. *Journal of Field Archaeology,* 33, 259–277.

Karavanić, I., Banda, M., Radović, S., Miko, S., Vukosavljević, N., Razum, I., Smith, F. H. (2021). A palaeoecological view of the last Neanderthals at the crossroads of south-central Europe and the central Mediterranean: long-term stability or pronounced environmental change with human responses. *Journal of Quaternary Science,* 1–10. Doi: 10.1002/jqs.3279

Katoh, K., Standley, D. M. (2013). MAFFT Multiple Sequence Alignment Software Version 7: Improvements in Performance and Usability. *Molecular Biology and Evoutionl,* 30, 772–780.

Klimowicz, M., Nadachowski, A., Lemanik, A., Socha, P. (2016). Is enamel differentiation quotient (SDQ) of the narrow-headed vole (*Microtus gregalis*) useful for the Pleistocene biostratigraphy? *Quaternary International,* 420, 348-356.

Koliushev, I. I. (1966). O zhyvotnom mirie pescher. In: Komendar, V.I., Artemhuk, I.V., Ivanov, S.D., Stoiko, S.M., Trybun, P.A., Turianin, I.I. (Eds.), Karpatskiye Zapovedniki. Karpaty, Uzhgorod, pp. 46–53.

Komšo, D. (2008). Limski kanal (Lim Channel). In: Matica B. (Ed.), Hrvatski arheološki godišnjak 4 (2007), 266–267. [in Croatian with English summary].

Krajcarz, M., Madeyska, T. (2010). Application of the weathering parameters of bones to stratigraphical interpretation of the sediments from two caves (Deszczowa Cave and Nietoperzowa Cave, Kraków-Częstochowa Upland, Poland). *Studia Quaternaria,* 27, 43-54.

Krochko, Y. I., Korchynskyi, O. V., Vargovych, R. S. (1993). Antropogenovi kistkovi zakhoronennia khrebetnykh tvaryn Zakarpattya. In: Fauna Skhidnykh Karpat: Suchasnyi Stan i Ohorona. Uzhgorod, pp. 84–85.

Kryštufek, B. (2008). *Glis glis*. Mammalian species, 195-206.

Lebreton, L., Soriano, S. (2018). Micromammals taphonomy and palaeoenvironmental reconstruction of a late Pleistocene site: a case study at Roc-en-Pail (France). *Quaternaire,* 29(1), 69-74.

Li et al. 2009

Lorenc, M. (2013). Radiocarbon ages of bones from Vistulian (Weichselian) cave deposits in Poland and their stratigraphy*. Acta Geologica Polonica,* 63(3), 399-424.

Ložek, V., Gáal, L., Holec, P., Horáček, I. (1989). Stratigraphy and Quaternary fauna of the cave Peskö in the Rimava basin. *Slovenský kras,* 27, 29-56.

Luzi, E., Berto, C., Calattini, M., Tessaro, C., Galiberti, A., Sala, B. (2020). Non-analogue rodent community in Central Italy during late Pleistocene: the case of Grotta del Sambuco, in: Proceedings of INQUA SEQS 2020 Conference, Wrocław, Poland. P. 70.

Malez, M. (1962). Romualdo Cave – a new significant Pleistocene site in Istria. *Bulletin Scientifique,* 7(6), 159–160.

Malez, M. (1968). Tragovi paleolita u Romualdovoj pećini kod Rovin­ja u Istri (Paläolitische Spuren in der Romualdohöhle bei Rovinj in Istrien). *Arheološki radovi i rasprave Jugoslavenske akademije znanosti i umjetnosti,* 6, 7–26. [in Croatian with German summary]

Malez, M. (1978). Paleontološka i kvartargeološka istraživanja u 1973. godini (Paleontological and geological investigation of the Quaternary in the year 1973). *Ljetopis Jugoslavenske akademije znanosti i umjet­nosti,* 78, 559–572. [in Croatian].

Malez, M., (1987). Pregled paleolitičkih i mezolitičkih kultura na području Istre (Übersicht der Paläolithischen und Mesolithischen Kulturen in Istrien*). Izdanja Hrvatskog arheološkog društva,* 11, 3–47. [in Croatian with German summary].

Maricic, T., Whitten, M., & Pääbo, S. (2010). Multiplexed DNA sequence capture of mitochondrial genomes using PCR products. *PLoS ONE*, 5, 9–13.

Marković, Z. (2008). Small mammals (Rodentia and Lagomorpha) from Gradašnica Cave (East Serbia). *Bulletin of Natural History Museum Belgrade*, 1, 65-77.

Marković, Z., Pavlović, G. (1991). Prvi rezultati istraživanja faune Vrelske pećine (Bela Palanka, Srbija*). Geološki anali Balkanskoga poluostrva*, 55, 221–230.

Mauch Lenardić, J. (2013). First record of *Dicrostonyx* (Rodentia, Mammalia) in the Late Pleistocene/Holocene sediments of Croatia. *Geologia Croatica,* 66(3), 183–189. Doi: 10.4154/gc.2013.15.

Mauch Lenardić, J. (2014). Bank vole *Myodes* (=*Clethrionomys*) *glareolus* (Schreber, 1780): Rare species in the Late Pleistocene fauna of Croatia. *Quaternary International,* 328–329, 167–178. Doi: 10.1016/j.quaint.2013.07.131.

Mauch Lenardić, J., Oros Sršen, A., Radović, S. (2018). Quaternary fauna of the Eastern Adriatic (Croatia) with the special review on the Late Pleistocene sites. *Quaternary International,* 494, 130–151. <https://doi.org/10.1016/j.quaint.2017.11.028>.

Meyer, M., & Kircher, M. (2010). Illumina sequencing library preparation for highly multiplexed target capture and sequencing. *Cold Spring Harbor Protocols*, *5*(6), t5448. doi: 10.1101/pdb.prot5448

Milne, I., Stephen, G., Bayer, M., Cock, P. J. A., Pritchard, L., Cardle, L., … Marshall, D. (2013). Using Tablet for visual exploration of second-generation sequencing data. *Briefings in Bioinformatics*, *14*(2), 193–202. doi: 10.1093/bib/bbs012

Miracle, P. (2005). Late Mousterian Subsistence and Cave Use in Dalmatia: the Zooarchaeology of Mujina Pećina, Croatia. *International Journal of Osteoarchaeology,* 15, 84–105.

Mirea, I.-C., Robu, M., Petculescu, A., Kenesz, M., Faur, L., Arghir, R., Tecsa, V., Timar – Gabor, A., Roban, R.-D., Panaiotu, C.-G., Sharifi, A., Pourmand, A., Codrea, V. A., Constantin, S. (2020). Last deglaciation flooding events in the South Carpathians as revealed by the study of cave deposits from Muierilor Cave, Romania*. Palaeogeography, Palaeoclimatology, Palaeoecology,* 562, 110084.

Nadachowski, A., Harrison, D. L., Szyndlar, Z., Tomek, T., Wolsan, M. (1993). Late Pleistocene vertebrate fauna from Obłazowa 2 (Carpathians, Poland): paleoecological reconstruction. *Acta Zoologica Cracoviensia,* 36(2), 281-290.

Nadachowski, A., Żarski, M., Urbanowski, M., Wojtal, P., Miękina, B., Lipecki, G., Ochman, K., Krawczyk, M., Jakubowski, G., Tomek, T. (2009). Late Pleistocene environment of the Częstochowa Upland (Poland) reconstructed on the basis of faunistic evidence from archaeological cave sites. Institute of Systematics and Evolution of Animals, Polish Academy of Sciences, Kraków, 112 pp.

Nicolăescu-Plopșor, C. S., Comșa, A., Nicolăescu-Plopșor, D. C., Bolomey, A. (1957). “Baia de Fier archeological site”. *Materiale si Cercetări Arheologice,* 3, 13–27.

Obuch, J. (2006). Fosílna fauna belane tundrovej (*Nyctea scandiaca*) v jaskyni Šarkanica [Fossil food of the Snowy Owl (*Nyctea scandiaca*) in the Šarkanica cave]. *Reussia,* 3(2) 160.

Păunescu, A., Rădulescu, C., Samson, P. (1982). “Découvertes du Paléolithique Inférieur en Roumanie.” *Travaux de L`Institut de Speologie “Emile Racovitza*” XXI, 53–62.

Petculescu, A. (2013). Paleoclimatic reconstructions, on the basis of small mammals, from carstic deposits. Ed. AGIR, pp. 1-186, Bucharest.

Petrov, B. (1992). Mammals of Yugoslavia (Insectivores and Rodents). - Natural History Museum, Suppl. 37, Belgrade, 186 pp.

Petrović, J. (1976). Jame i pećine SR Srbije. - Vojna biblioteka. Pravila i udžbenici 39, Vojno izdavački zavod, Beograd. 551 pp. [in Serbian].

Popov, V. V. (1994). Quaternary small mammals from deposits in Temnata - Prochodna Cave system. In J. K. Kozlowski, H. Laville, and B. Ginter (Eds.), Temnata Cave, Excavations in Karlukovo Karst Area, Bulgaria (vol. 1, part 2, pp. 11 - 53). Krakow: Jagellonian University Press.

Popov, V. V. (2000). The small mammals (Mammalia: Insectivora, Chiroptera, Lagomorpha, Rodentia) from Cave 16 (North Bulgaria) and the paleoenvironmental changes during the Late Pleistocene. - In: B. Ginter, J. K. Kozlowski, K. Laville (eds.) Temnata Cave. Excavations in Karlukovo Karst Area, Bulgaria, (vol.2, part 1, pp. 13 - 65). Krakow, Jagellonian University Press.

Popov, V., Di Canzio, E., Giaccio, B. (2014). Late Quaternary Small Mammals and Paleotemperatures in Bulgaria and Italy. *Acta Zoologica Bulgarica*, 66, 89-108.

Popov, V. V. (2018). Pliocene-Quaternary Small Mammals (Eulipotyphla, Chiroptera, Lagomorpha, Rodentia) in Bulgaria: Biostratigraphy, Paleoecology, and Evolution. In: The Pleistocene: Geography, Geology, and Fauna (pp. 109 - 235). Editors: Gaetan Huard and Jeannine Gareau. Nova Science Publishers. ISBN: 978-1-53613-728-6; 978-1-53613-729-3 (eBook).

Quinlan, A. R., Hall, I. M. (2010). BEDTools: A flexible suite of utilities for comparing genomic features. Bioinformatics, 26, 841–842.

Reimer, P. J., Austin, W. E. N., Bard, E., Bayliss, A., Blackwell, P. G., Bronk Ramsey, C., et al. (2020). The IntCal20 Northern Hemisphere Radiocarbon Age Calibration Curve (0–55 cal kBP). *Radiocarbon,* 62, 725–757.

Richard, M., Falguères, C., Valladas, H., Ghaleb, B., Pons-Branchu, E., Mercier, N., Richter, D., Conard, N. J. (2019). New electron spin resonance (ESR) ages from Geißenklösterle Cave: A chronological study of the Middle and early Upper Paleolithic layers. *J. Hum. Evol*, 133, 133-145.

Ridush, B. (2012). Palaeogeographic records in sediments of karst caves in Ukrainian Carpathians. *Scientific Annals of Stefan cel Mare University of Suceava. Geography Series* 21, 80. doi:10.4316/georeview.2012.21.1.57

Ridush, B. (2014). “Bear Caves” of South-Eastern Europe. *Speleology and Karstology,* 12, 26–41.

Ridush, B., Popiuk, Y. (2019). New spicies from the bone accumulatioin Perlyna Cave (Ukrainian Carpathians). In: Kavcik-Graumann, N. (Ed.), 25th International Cave Bear Symposium. Universität Wien - National Park Paklenica, Paclenica, p. 8.

Rink, W. J., Karavanić, I., Pettitt, P. B., van der Plicht. J., Smith, F. H., Bartoll, J. (2002). ESR and AMS-based ^14^C dating of Mousterian levels at Mujina Pećina, Dalmatia, Croatia. *Journal of Archaeological Science,* 29, 943–952.

Ruiz-Redondo, A., Komšo, D., Garate Maidagan, D., Moro-Abadía, O., Gonzáles-Morales, M. R., Jaubert, J., Karavanić, I. (2019). Expanding the horizons of Palaeolithic rock art: the site of Romualdova Pećina. *Antiquity* 93, 368, 297–312.

Sekyra, J. (1954). Velehorský kras Bělských Tater. Nakl. ČSAV, Praha.

Schaefer, H. (1975). Die Spitzmäuse der Hohen Tatra seit 30000 Jahren (Mandibular Studie). *Zoologischen Anzeiger*, 195(1/2), 89-111.

Schubert, M., Lindgreen, S., & Orlando, L. (2016). AdapterRemoval v2: Rapid adapter trimming, identification, and read merging. *BMC Research Notes*, *9*(1), 1–7. doi: 10.1186/s13104-016-1900-2

Shapiro, B., Ho, S. Y. W., Drummond, A. J., Suchard, M. A., Pybus, O. G., Rambaut, A. (2011). A bayesian phylogenetic method to estimate unknown sequence ages. *Molecular Biology and Evolution,* 28, 879–887.

Soficaru, A., Doboș, A., Trinkaus, E. (2006). “Early Modern Humans from the Pestera Muierii, Baia de Fier, Romania. *Proceedings of the National Academy of Sciences of the United States of America,* 103(46), 17196–201.

Soriano, S., Ahmed-Delacroix, N., Borvon, A., Chevrier, B., David, E., Dessoles, M., Voeltzel, B. (2021). Le site paléolithique de Roc-en-Pail (Chalonnes-sur-Loire, Maine-et-Loire). État des connaissances 150 ans après sa découverte. *Gallia Préhistoire*, 61. https://journals.openedition.org/galliap/2633.

Stojak, J., McDevitt, A. D., Herman, J. S., Searle, J. B., Wójcik, J. M. (2015). Post-glacial colonization of eastern Europe from the Carpathian refugium: evidence from mitochondrial DNA of the common vole Microtus arvalis. *Biological Journal of the Linnean Society*, 115, 927–939.

Suárez-Bilbao, A., Elorza, M., Castaños, J., Arrizabalaga, A., Iriarte-Chiapusso, M. J., Murelaga, X. (2018). The Late Pleistocene avifauna from Artazu VII (Basque Country, northern Iberian Peninsula). *Historical Biology*, 32(3), 307–320.

Suárez-Bilbao, A., Garcia-Ibaibarriaga, N., Arrizabalaga, A., IriarteChiapusso, M. J., Murelaga, X. (2017). Paleoenvironmental and Paleoclimatic approach to the late Pleistocene site of Artazu VII (Arrasate, northern Iberian Peninsula) using small mammals. *Ameghiniana*, 54(6), 641–654.

Suárez-Bilbao, A., García-Ibaibarriaga, N., Castaños, J., Castaños, P., Iriarte, M.J., Arrizabalaga, A., Torres, T., Ortiz, J.E., Murelaga, X. (2016). A new Late Pleistocene non-anthropogenic vertebrate assemblage from the northern Iberian Peninsula: Artazu VII (Arrasate, Basque Country). *C. R. Palevol,* 15, 950-957.

Tatarinov, K. A., Bachynskyi, G. A. (1968). Peshchernyje zakhoronenia pliotsenovykh i antropogenovykh pozvonochnykh v zapadnykh obllastiakh Ukrainy [Cave burials of Pliocene and Anthropogene Vertebrates in the western regions of Ukraine]. *Bulletin of the Moscow Society of Naturalist,* 73, 114–121.

Terzea, E. (1971). Les Micromammiferes quaternaires de deux grottes des Carpates roumaines. *Travaux de L`Institut de Speologie “Emile Racovitza”* X, 279–300.

Terzea, E. (1986). Chronologie des faunes Pleistocenes superieures du sud-ouest de la Roumanie. *Travaux de L`Institut de Speologie “Emile Racovitza”* XXV, 85–101.

Utge, J., Sévêque, N., Lartigot-Campin, A. S., Testu, A., Moigne, A. M., Vézian, R., Elalouf, J. M. (2020). A mobile laboratory for ancient DNA analysis. *Plos One,* 15(3), e0230496.

van Klinken, G. J. (1999). Bone collagen quality indicators for palaeodietary and radiocarbon measurements. *Journal of Archaeological Science,* 26(6), 687–695.

Wacker, L., Němec, M., Bourquin, J. (2010a). A revolutionary graphitisation system: Fully automated, compact and simple, Nuclear Instruments and Methods in Physics Research, Section B: Beam Interactions with Materials and Atoms, 268, 931–934.

Wacker, L., Bonani, G., Friedrich, M., Hajdas, I., Kromer, B., Němec, M., Ruff, M., Suter, M., Synal, H. A., Vockenhuber, C. (2010b). Micadas: Routine and high-precision radiocarbon dating, *Radiocarbon,* 52, 252–262.

Wacker, L., Fahrni, S. M., Hajdas, I., Molnar, M., Synal, H. A., Szidat, S., Zhang, Y. L. (2013). A versatile gas interface for routine radiocarbon analysis with a gas ion source, Nuclear Instruments and Methods in Physics Research, Section B: Beam Interactions with Materials and Atoms, 294, 315–319.

Wolf, S. (2013). Schmuckstücke - Die Elfenbeinbearbeitung im Schwäbischen Aurignacien. Eberhard-Karls Universität Tübingen, Tübingen.
